# Cap-specific second nucleotide ribose methylase CMTR2 is required for transcriptome transition during mammalian germline development

**DOI:** 10.64898/2026.07.28.741271

**Authors:** Harsha Raheja, Elena Delfino, Michaela Dohnalkova, Richard J. Fish, Cathrine Broberg Vågbø, Fabienne Burger, Carmen Fernandez-Rodriguez, David Homolka, Ramesh S. Pillai

## Abstract

Eukaryotic RNA polymerase II transcripts carry a signature m7G cap (Cap0) structure that is co-transcriptionally added to the 5’ end, and is essential for translation and RNA stability. Higher eukaryotes carry additional essential ribose methylations on the first and second cap-proximal nucleotides, termed as Cap1 and Cap2, respectively. The ubiquitous Cap1 modification protects cellular RNAs from being recognized by the innate immune sensors. Cap2 is also implicated in such an innate immune role, but here we use our genetic analyses of two human cell lines and three mouse tissues to reveal that loss of CMTR2 does not result in activation of the innate immune response. Germline deletion of mouse CMTR2 shows that it is required for male and female fertility. While mutant germ cells proceed into the meiotic pachytene spermatocyte stage, their transcriptome fails to keep pace and transition from the preceding leptotene/zygotene stages. Such a meiotic role is not conserved in other vertebrates like zebrafish, as *cmtr2* mutants are fertile, instead it has a role in defining sex, as all mutants are exclusively males. Taken together, our study reveals that CMTR2 does not influence innate immune response in human cells and mouse tissues, but shapes gene expression during developmental transitions.

## INTRODUCTION

The *N^7^*-methyl guanosine (m7G) cap structure is co-transcriptionally added to RNA polymerase II transcripts via a 5’-5’ triphosphate bridge (Cowling, 2009; Galloway & Cowling, 2019; Pelletier *et al*, 2021). Termed Cap0 structure, this modification is found in all eukaryotes and essential for translation and RNA stability. Additional methylations were discovered on cap-proximal nucleotides in human polyA+ RNA (Keith *et al*, 1978; Wei *et al*, 1975a; Wei *et al*, 1975b). The transcription start site (TSS) nucleotide is methylated on the ribose (Nm) by CMTR1 (Belanger *et al*, 2010), creating the Cap1 structure (Figure 1A), while the second nucleotide is ribose methylated by CMTR2 to create the Cap2 structure (Werner *et al*, 2011). When the TSS of Cap1 RNAs is an adenosine, it is further *N^6^*-methylated (m^6^A) by PCIF1 (Akichika *et al*, 2019), creating the m^6^Am modification. The Cap1 modification is established as a discriminator of self vs non-self RNAs (Dohnalkova *et al*, 2023; Schoggins *et al*, 2011; Schuberth-Wagner *et al*, 2015), as it reduces affinity of innate immune sensors for cellular RNAs (Kowalinski *et al*, 2011), preventing innate immune response activation. Cytosolic viruses encode their own Cap1 methylase to mark viral RNAs as a mechanism to evade the host innate immune system (Daffis *et al*, 2010; Zust *et al*, 2011).

**Figure 1.**
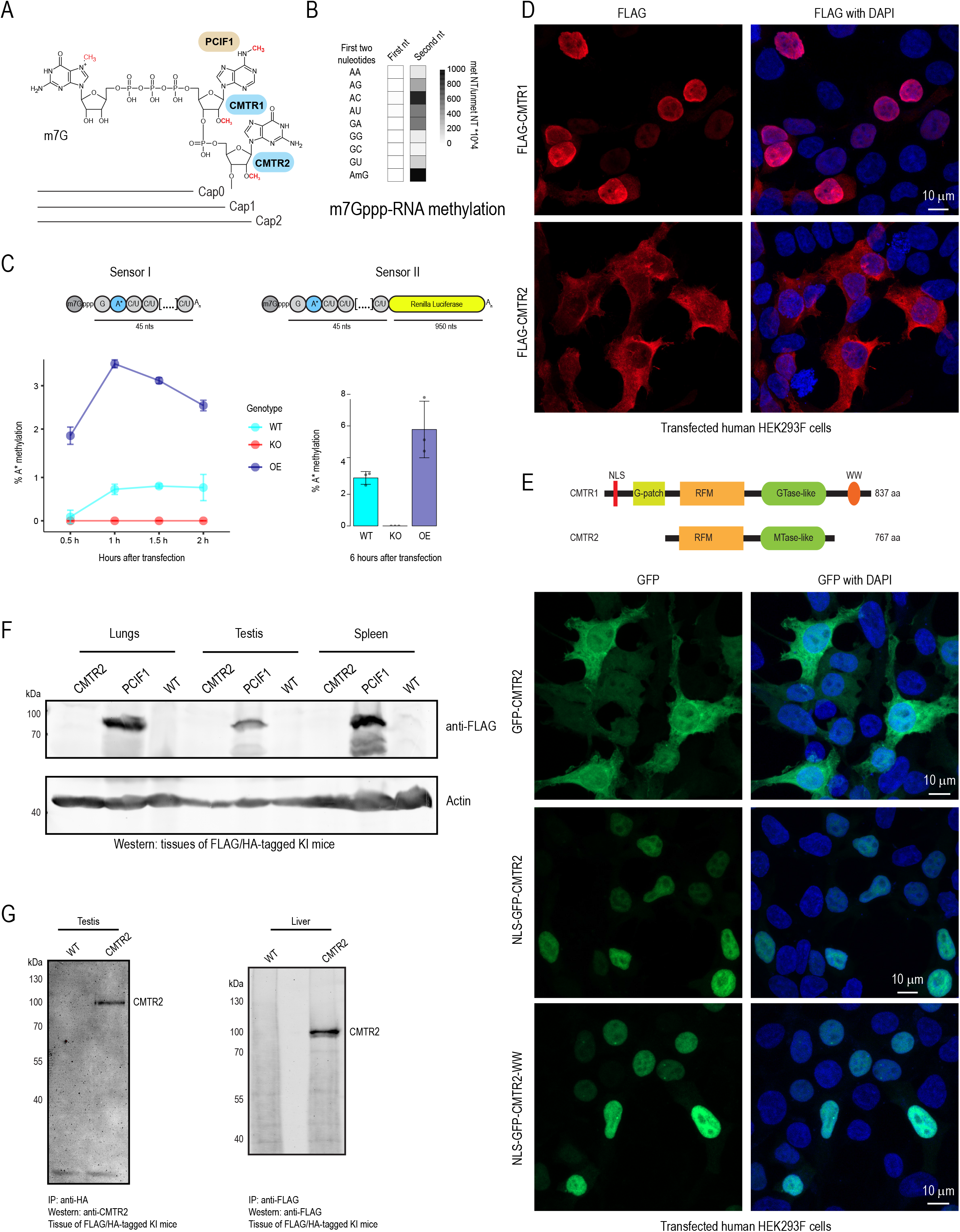
CMTR2 is a cytoplasmic low abundant Cap2 methylase. (A) Schematic depicting the modifications on the 5’ cap proximal nucleotides of mRNAs from higher eukaryotes. The enzymes that catalyze the depicted modifications are mentioned beside the modifications. (B) Recombinant mouse CMTR2 was used to *in vitro* methylate capped RNAs. RNA substrates with the indicated first and second nucleotides were analyzed for methylation at the corresponding cap-proximal position. The indicated nucleotides were unique and not present elsewhere in the body of the transcript. The number of methylated nucleotides per 10,000 of the respective unmethylated nucleotide is shown. (C) RNA sensors were designed to quantify 2’-O-methylation of the second nucleotide. Schematics illustrate the nucleotide composition of the sensors. The second nucleotide (blue-A*) was heavier than the regular nucleotides (composed of ^15^N, ^13^C in place of ^14^N and ^12^C) to be differentiated by LC-MS. SensorI, a non-coding RNA, was transfected to HEK293F wildtype (WT), CMTR2 Knockout (KO) or CMTR2 overexpression (OE) cells and analysed at the indicated time points (left). SensorII, which encodes Renilla luciferase and contains the same heavy second nucleotide, was transfected into WT, KO and OE cells for 6 h (right). At the indicated time points, cells were harvested, poly (A+) RNA purified and the methylation status of heavy second nucleotide determined by LC-MS. The percentage of methylated heavy second nucleotide (A*) is shown. n=3 (D) Subcellular localization of cap ribose methylases. 3XFLAG-HA-CMTR1 and 3XFLAG-HA-CMTR2 expressing plasmids were transfected to HEK293F cells. The cells were harvested 24 h after transfection and the localization of CMTR1/2 detected by immunofluorescence staining using anti-FLAG primary antibody and AF647-conjugated secondary antibody (Red). Nuclei were counterstained with DAPI (Blue). Scale bar represents 10 μm. (E) Alteration of CMTR2 subcellular localization. The schematic depicts domain organization of CMTR1 and CMTR2 proteins. NLS and WW domains from CMTR1 were cloned in GFP-CMTR2 expressing plasmids in the order mentioned alongside the images. Plasmids expressing the indicated GFP fusion proteins were transfected into HEK293F cells. Cells were imaged 24 h after transfection for GFP expression analysis (Green). Nuclei were counterstained with DAPI (Blue). Scale bar represents 10 μm. (F) Tissue expression analysis of *3XFLAG-HA-Cmtr2* (CMTR2) knock-in mouse. *Pcif1-3XFLAG* (PCIF1) mouse tissues were used as positive control and wild type (WT) mouse tissues were used as negative control. Western blotting was performed using anti-FLAG antibody. β-actin was used as the loading control. (F) Immunoprecipitation of endogenous CMTR2 from liver and testis of *3XFLAG-HA-Cmtr2* knock-in mice. Liver and testicular lysates from *3XFLAG-HA-Cmtr2* (CMTR2) knock-in mouse were immunoprecipitated using anti-FLAG /anti-HA beads. Tissues from WT mouse were used as control. Western blotting was performed using anti-CMTR2/ anti-FLAG antibodies as indicated.

Cap1 or m6Am is found to be ubiquitously present on human polyA+ RNAs (Akichika *et al*., 2019; Dix *et al*, 2022). On the other hand, a Cap2 quantification and mapping protocol has established that the modification is present on a subset of polyA+ RNAs (Despic & Jaffrey, 2023). The same study used a *CMTR2* knockout (KO) human HEK293T cells to reveal that the innate immune pathway is activated in the mutant cells (Despic & Jaffrey, 2023). Cap2 was found not to affect RNA stability or translation, while another study using transfected mRNAs, demonstrated an influence of Cap2 on translation, dependent on the cell line used (Drazkowska *et al*, 2022). *Cmtr2* is not essential in worms, but when combined with *Cmtr1* deletion, accentuates the consequences for post-embryonic growth and fertility (Clemens *et al*, 2026). In flies, *Cmtr1* and *Cmtr2*, and their double-mutants are viable, but display learning defects (Haussmann *et al*, 2022). Fly ribose methylases are shown to act redundantly as a Cap1 methylase (Haussmann *et al*., 2022). In fact, Cap2 modification is barely detected in polyA+ RNAs from worms and very low in flies (Clemens *et al*., 2026; Despic & Jaffrey, 2023).

We (Dohnalkova *et al*., 2023) and others (Yermalovich *et al*, 2024), previously showed that loss of mouse *Cmtr2* results in embryonic lethality, but without activating the interferon pathway, as the innate immune sensors are not yet expressed during the mid-gastrulation developmental window where the arrest happens. Thus, there are physiological and gene expression consequences of losing Cap2 methylation, which remain to be discovered. Here, we used human *CMTR2* KO cell lines and mouse conditional deletion in liver, lung and the germline, to find that there is no activation of the innate immune pathway. Instead, deletion of *Cmtr2* in the germline results in mouse male and female sterility. Transcriptome analysis reveals that arrest in meiotic progression in the *Cmtr2* mutant male germline is due to a failure to transition the transcriptome during germ cell development. While this meiotic role is not conserved in other vertebrates like zebrafish, we show that zebrafish *cmtr2* is required for defining the sex, as all mutants are exclusively males.

## RESULTS

### Cap2 sensor RNA reveals that nucleo-cytoplasmic CMTR2 is extremely low abundant and can act post-transcriptionally on targets

Since *Drosophila* (Haussmann *et al*., 2022) and *C. elegans* (Clemens *et al*., 2026) *Cmtr2* act redundantly with the Cap1 methylase *Cmtr1*, we wished to directly investigate the biochemical activity of mouse CMTR2 (Figure 1A). We produced the recombinant mouse CMTR2 (Figure S1A) and incubated it with several m7G-capped RNAs (Cap0) (Figure S1G). The reaction products were subject to RNA mass spectrometry. In all the nine tested RNAs, mouse CMTR2 exclusively methylates the second nucleotide, confirming it to be a Cap2 methylase (Figure 1B and Figure S1G).

To monitor CMTR2-mediated Cap2 methylation within cells, we designed a sensor RNA (Figure 1C; sensor I). Briefly, it is a short m7G-capped and polyadenylated noncoding RNA which has a second nucleotide that is labelled with heavy isotopes (resulting in it being 15 Daltons heavier than the endogenous counterpart). After transfection into human HEK293F cells, polyA+ RNA was isolated at 30 min intervals and subject to RNA mass spectrometry. The heavy isotope-labelled nucleoside (Figure S1H) and its methylated version was specifically detected and quantified (Figure 1C). Cap2 methylation was detected in wildtype (WT) cells as early as 30 mins post-transfection and increased at later time points. This methylation was boosted upon CMTR2 overexpression (OE). Methylation was not detected in CMTR2 knockout (KO) cells, confirming the specific activity of endogenous CMTR2 for Cap2 methylation (Figure 1C). Next, we generated a coding sensor RNA, which also has heavy adenosine at the second nucleotide (Figure 1C and Figure S1H; sensor II). Cap2 methylation was detected in WT HEK293F cells, but not in the CMTR2 KO. Similar increase in sensor Cap2 methylation upon CMTR2 overexpression confirmed the low basal abundance of endogenous CMTR2. We confirmed luciferase expression from the transfected RNA. Furthermore, this also shows that CMTR2 acts on any available m7G-capped RNAs, independently of the translation status of the RNA.

To verify their site of action within the cell, we tagged the two ribose methylases with a 3xFLAG-HA tag and transiently expressed the fusions in human cells. Immunofluorescence analysis indicates that CMTR1 is exclusively nuclear and shows a diffused distribution in the nucleoplasm. In contrast, CMTR2 is diffusely present in both the cytoplasm and nucleus (Figure 1D). CMTR1 has a nuclear localization signal (NLS), lacking in CMTR2 (Figure 1E). We made GFP-tagged CMTR2 expression constructs where we inserted the NLS from CMTR1, and also with an additional WW domain that allows it to interact with the C-terminal domain (CTD) of RNA polymerase II (pol II). Localization studies show that CMTR2 fusion proteins artificially tagged with the NLS and WW domain are exclusively nuclear (Figure 1E), like CMTR1. These studies show that CMTR2 lacks strong nuclear targeting signals which makes it nucleo-cytoplasmic in human cells, and likely accesses RNAs post-transcriptionally.

The low level of endogenous CMTR2 activity and its increase in over-expression conditions (Figure 1C), indicated that CMTR2 is likely maintained at very low levels in vivo. To verify this, we created a knock-in mouse line where CMTR2 is endogenously tagged with N-terminal FLAG-HA tags (Figure S1C-D). Homozygous knock-in animals are viable, indicating that the tagged protein is functional, as loss of *Cmtr2* leads to embryonic lethality (Dohnalkova *et al*., 2023). The tagged CMTR2 protein is non-detectable in multiple mouse tissue lysates (Figure 1F and Figure S1E), and was only faintly visible after enrichment via immunoprecipitation (Figure 1G and Figure S1F). In contrast, expression of FLAG-HA-tagged PCIF1, another cap-specific methylase (Figure 1A), is abundantly detected in mouse tissue lysates, and did not need enrichment for visualization (Figure 1F and Figure S1E). Taken together, these results indicate that CMTR2 is a nucleo-cytoplasmic Cap2 methylase that is extremely low abundant in vivo. We further show that it likely acts on any accessible targets post-transcriptionally, and without any relevance to its translation status.

### Loss of CMTR2 in human cells and mouse tissues does not activate the innate immune pathway

To investigate the role of CMTR2 in vivo, we generated one clone of *CMTR2* KO human embryonic kidney 293F cells (HEK293F) and two clones of *CMTR2* KO A549 lung carcinoma cells. Western analysis confirms the absence of the protein in the two KO cell lines (Figure 2A). Transcriptome analysis indicates that ∼10-15% of the genes were either up or down in the mutants, when compared to the control cells (Figure 2B). Control cells went through the same procedure for mutagenesis but were selected for absence of edits in the target locus. Thousands of transcripts are dysregulated in the three cell lines (Figure 2C-E). The two A549 CMTR2 KO cell lines have largely similar expression profile (Figure S2A), with over 2500 dysregulated genes shared between them (Figure S2C). A previous study using HEK293T cells reported an activation of the cellular innate immune pathway (Despic & Jaffrey, 2023), but we did not observe any broad activation of the Type I interferon-mediated signalling pathway genes in the HEK293F CMTR2 KO mutant cell line (Figure 2F) and did not find any related GO terms enriched among upregulated genes (Figure S2C). This was also true for the A549 CMTR2 KO #1, with KO#2 showing only a few genes being dysregulated (Figure 2F). Taken together, our three CMTR2 KO human cell line models do not support a general role for CMTR2 in suppression of the innate immune response pathway. It is possible that levels of innate immune sensors and genetic compensatory pathways that arise during cell line establishment, might influence differences we observed in our three cell lines, and with that published previously (Despic & Jaffrey, 2023).

**Figure 2.**
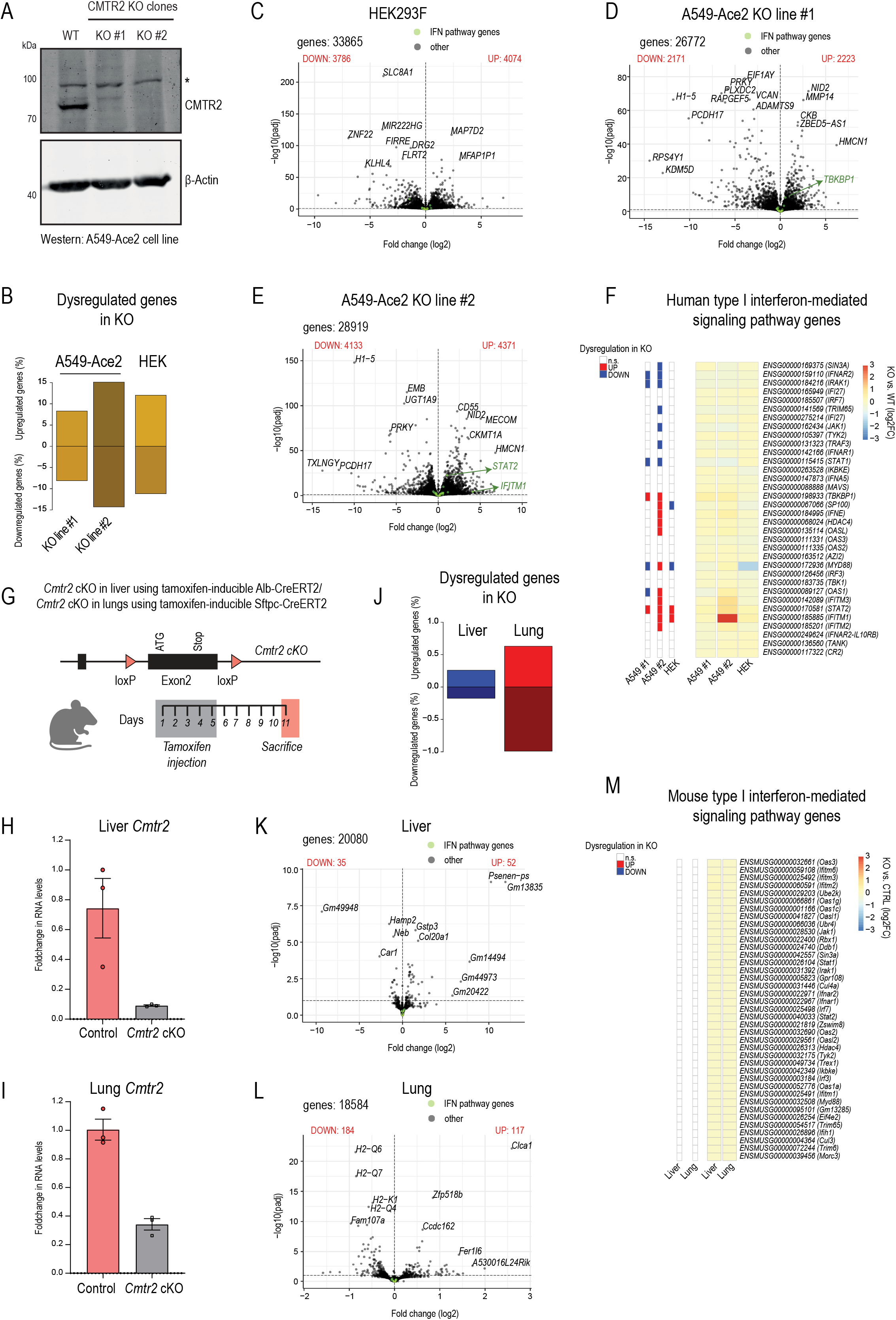
Loss of Cmtr2 is not associated with innate immune activation. (A) Confirmation of A549-Ace2 knockout clones. Western blotting was performed for wildtype (WT), CMTR2 knockout clone 1 (KO#1) and CMTR2 knockout clone 2 (KO#2) cells using anti-CMTR2 antibody. β-actin was used as the loading control. (B) Plot for the percentage of upregulated and downregulated genes in CMTR2 KO cell lines as compared to the respective WT cell line. (C) Volcano plot showing differential gene expression profile of the CMTR2 KO HEK293F cell line as compared to the WT HEK293F cell line. The genes associated with Interferon pathway are highlighted in green. (D) Volcano plot showing differential gene expression profile of the CMTR2 KO A549-Ace2 cell line 1 (KO#1) as compared to the WT A549-Ace2 cell line. The genes associated with Interferon pathway are highlighted in green. (E) Volcano plot showing differential gene expression profile of the CMTR2 KO A549-Ace2 cell line 2 (KO#2) as compared to the WT A549-Ace2 cell line. The genes associated with Interferon pathway are highlighted in green. (F) Heatmap showing the expression pattern of Human type I interferon-mediated signalling pathway genes in CMTR2 KO cell lines as compared to the respective WT cell line (right). Differentially expressed genes (adjusted p-value < 0.1) are highlighted in red (upregulated) or blue (downregulated) (left). (G) Strategy used for the generation of *Cmtr2* conditional knockout (cKO) in mouse liver and lung by deleting Exon 2 using tamoxifen inducible liver specific (Alb-CreERT2) or lung specific (Sftpc-CreERT2) cre. (H) *Cmtr2* RNA levels in Control and *Cmtr2* cKO liver were quantified by real-time PCR using gene specific primers. Fold change in the RNA levels was quantified by normalization to actin RNA levels. n=3; error bars refer to standard deviation. (I) *Cmtr2* RNA levels in Control and *Cmtr2* cKO lungs were quantified by real-time PCR using gene specific primers. Fold change in the RNA levels was quantified by normalization to actin RNA levels. n=3; error bars refer to standard deviation. (J) Plot for the percentage of upregulated and downregulated genes in *Cmtr2* cKO liver (*Cmtr2^-/loxP^; Alb-CreERT2^+/KI^*) / lung (*Cmtr2^-/loxP^; Sftpc-CreERT2^+/KI^*) as compared to the respective Control tissues (*Cmtr2^+/loxP^; Alb-CreERT2^+/KI^ / Cmtr2^+/loxP^; Sftpc-CreERT2^+/KI^*). (K) Volcano plot showing differential gene expression profile of the *Cmtr2* cKO liver as compared to the Control liver. The genes associated with Interferon pathway are highlighted in green. (L) Volcano plot showing differential gene expression profile of the *Cmtr2* cKO lung as compared to the Control liver. The genes associated with Interferon pathway are highlighted in green. (M) Heatmap showing the expression changes of Mouse type I interferon-mediated signalling pathway genes in *Cmtr2* cKO tissues as compared to the respective Control tissues (right). None of the genes are differentially expressed (adjusted p-value < 0.1) (left).

Cap-proximal ribose methylation on the transcription start site nucleotide (Cap1 modification) is a mechanism to mark cellular RNAs as ‘self’ to prevent their recognition by cellular innate immune sensors. We previously showed that deletion of the Cap1 methylase *Cmtr1*, in mouse liver displayed chronic activation of the innate immune response pathway (Dohnalkova *et al*., 2023). To test the relevance of Cap2 methylation using an inducible deletion strategy and one that does not involve long term adaptation, we obtained a floxed allele of *Cmtr2*, that allows conditional deletion of the gene in selected tissues (Figure 2G and Figure S3A). We prepared mice where the only functional copy of the *Cmtr2* allele is floxed with loxP sites (*Cmtr2 ^loxP/-^*) (Figure S3B-C). Crossing it with mice expressing CreERT2 recombinase, allowed tissue-specific inducible deletion of *Cmtr2*.

We used the *Albumin*-CreERT2 line to delete *Cmtr2* in mouse liver and the Sftpc-CreERT2 line to delete the gene in the lung tissue (Figure S3B-C). Loss of *Cmtr2* expression in the conditional deletion tissues was confirmed by quantitative reverse-transcription PCR (qRT-PCR) (Figure 2H-I). Histological examination of the mutant tissues did not reveal any major compositional changes (Figure S3E). Transcriptome analyses indicate minimal dysregulation in either tissue, with more alterations seen in the lung (Figure 2J). Within each tissue, up to 100 to 300 genes are altered (Figure 2K-L). Individual examination of the Type I interferon-stimulated signalling pathway genes shows that none are altered in both tissues (Figure 2M). Taken together, our analysis of human CMTR2 knockout cell lines and mutant mouse tissues shows that CMTR2, and by that extension Cap2 methylation, may not have a major role in suppression of the innate immune pathway as performed by Cap1 modification.

### Loss of CMTR2 in the mammalian germline causes male and female infertility

To test the relevance of *Cmtr2* in an environment that undergoes developmental changes, we deleted the gene using the germline-specific Ddx4 (Vasa)-Cre, whose expression commences at E14.5 in both male and female germlines (Figure 3A and Figure S3D). Female conditional knockout (cKO) animals were sterile (Figure 3B), with only rare births (1 pup in one litter) in crosses with wildtype partners. To investigate the impact of loss of *Cmtr2*, we performed histological analysis of adult (post-natal day 75, P75) ovaries (Figure 3C). Female germ cells enter meiosis at E13.5, prior to deletion of *Cmtr2*. By birth, the oocytes are arrested in meiosis I and held within primordial follicles. Observation of adult ovaries from control and germline cKO *Cmtr2* females shows that both have primary follicles with oocytes held in a single layer of somatic granulosa cells (Figure 3C). Further folliculogenesis is evident in the cKO, as secondary follicles with two layers of surrounding granulosa cells or more mature pre-antral follicles with fluid-filled cavities are visible (Figure 3C). The final stage antral or ovulatory follicles are also visible in both control and the cKO female ovaries. Thus, while folliculogenesis appears to be normal, females lacking *Cmtr2* in the germline are sterile.

**Figure 3.**
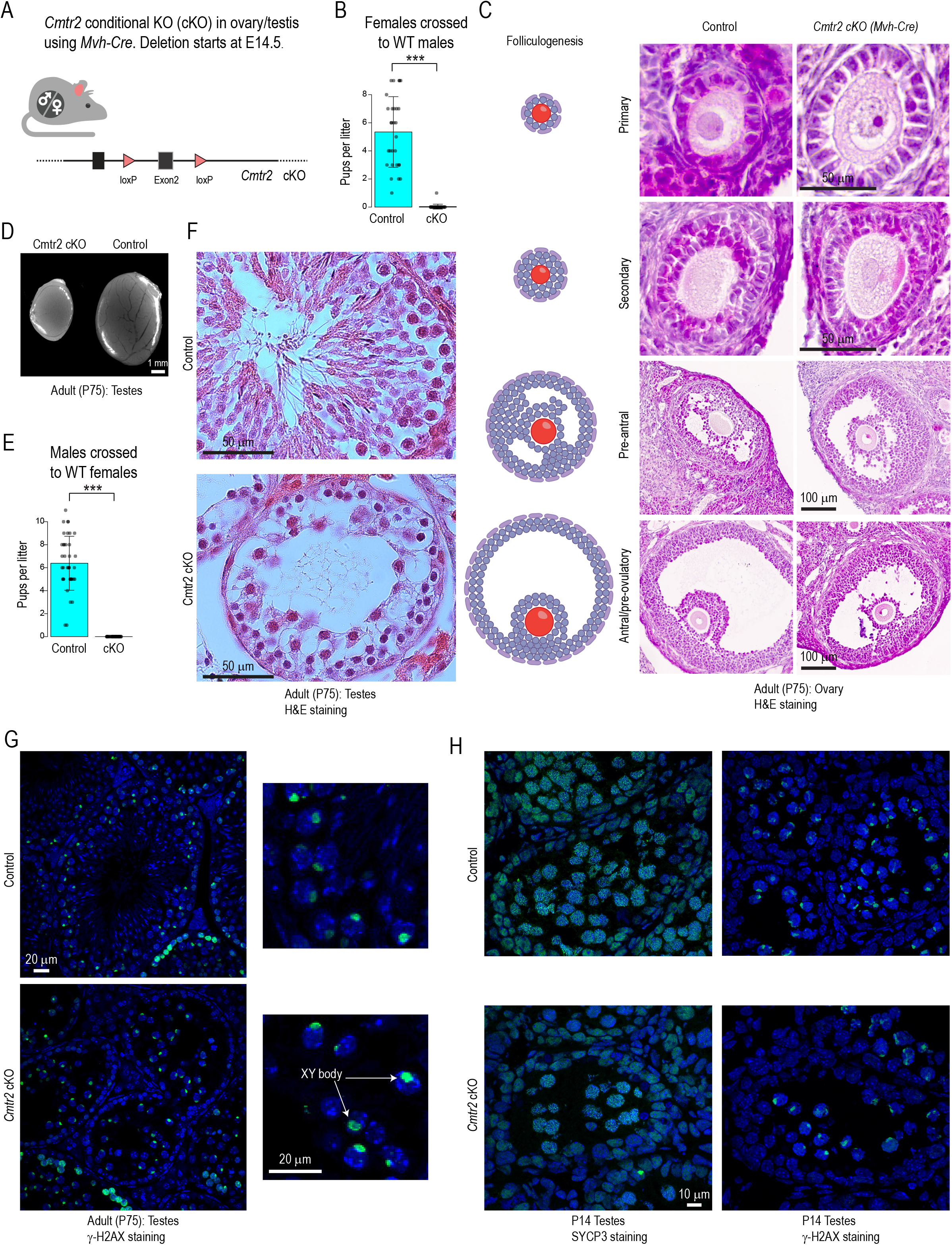
Mouse *Cmtr2* is essential for germline development and fertility. (A) Strategy used for the generation of *Cmtr2* conditional knockout (cKO) in mouse germline by deleting Exon 2 using DDX4 (Mvh)-Cre. (B) Fertility analysis of *Cmtr2* cKO females. Control (*Cmtr2^+/loxP^; DDX4-Cre^+/KI^)* and *Cmtr2* cKO (*Cmtr2^-/loxP^; DDX4-Cre^+/KI^*) females were bred with WT males and the number of pups born per litter analyzed. Error bars refer to standard deviation; *** p-value < 0.001 (t.test). (C) Histological analysis of hematoxylin and eosin-stained sections of Control and *Cmtr2* cKO ovaries. The cartoon represents different stages of follicular development. Scale bar represents 50/100 μm as indicated. (D) Atrophied testes upon *Cmtr2* cKO. Comparison of Control and *Cmtr2* cKO testes from adult animals (P75). Scale bar represents 1 mm. (E) Fertility analysis of *Cmtr2* cKO males. Control and *Cmtr2* cKO males were bred with WT females and pups born per litter analyzed. Error bars refer to standard deviation; *** p-value < 0.001 (t.test). (F) Histological analysis of hematoxylin and eosin-stained sections of Control and *Cmtr2* cKO testes (P75). Scale bar represents 50 μm. (G) γ-H2AX staining in Control and *Cmtr2* KO testes (P75). The testicular sections from Control and *Cmtr2* cKO animals were used for immunofluorescence staining using anti-γ-H2AX primary antibody and AF488-conjugated secondary antibody (Green). Nuclei were counterstained with DAPI (Blue). Scale bar represents 20 μm. (H) SYCP3 and γ-H2AX staining in Control and *Cmtr2* KO testes (P14). The testicular sections from Control and *Cmtr2* cKO animals were used for immunofluorescence staining using anti-SYCP3 and anti-γ-H2AX primary antibody and AF488-conjugated secondary antibody (Green). Nuclei were counterstained with DAPI (Blue). Scale bar represents 10 μm.

Males with cKO of *Cmtr2* in the germline display atrophied testes (Figure 3D) and were sterile (Figure 3E). Histological examination of adult testes shows that control animals had germ cells in all stages of meiosis with pachytene spermatocytes and post-meiotic haploid round spermatids and elongate spermatids and mature sperm (Figure 3F). In contrast, cKO animals have narrower tubules, fewer germ cells, and lack post-meiotic round spermatids, elongate spermatids and sperm (Figure 3F), explaining the atrophied testes in the mutant (Figure 3D). Males initiate meiosis after birth at around P8 to enter the prophase I, when leptotene spermatocytes are visible in the seminiferous tubules. To facilitate meiotic DNA recombination between the homologous chromosomes from the two parents, double-stranded breaks are generated in the leptotene stage. The homologous chromosomes pair by lying side-by side and this is facilitated by the synaptonemal complex. SYCP3 is a component of this complex that assembles along the chromosomes to form the lateral axes in the zygotene stage. Immunofluorescence analysis for SYCP3 shows that the protein stains along the length of the chromosomes in germ cells from both the control and *Cmtr2* cKO P14 testes (Figure 3H). At pachytene stage when the synapsis is complete, DNA crossovers are completed between homologous autosomes. While the synapsed autosomes remain transcriptionally active, the sex chromosomes that lack homology remain largely unsynapsed and are sequestered into a transcriptionally silent domain called the XY body or sex body. Several epigenetic marks are localized to this structure, including the phosphorylated histone H2AX (γ-H2AX), which is observed in both the control and cKO germ cells, indicating that germ cells proceed to the pachytene stage in the absence of *Cmtr2* (Figure 3G-H). Taken together, our genetic and histological analyses show that *Cmtr2* is required for fertility of both sexes and in males the mutant germ cells arrest at the pachytene stage of meiosis.

### CMTR2 regulates meiotic progression by facilitating transition of transcriptomes

To investigate the gene expression changes in the *Cmtr2* mutant male germ cells, we used FACS to purify meiotic germ cells with duplicated DNA (4n) (STAR Methods). We used control and *Cmtr2* cKO males at P9 (after initiation of meiosis) and at P14 (when germ cells are at the pachytene stage). Histological analysis shows uniform germ cells populations in the testes in both genotypes (Figure S4A-B). Germ cells of both genotypes display the γ-H2AX-labelled sex body in the P14 testes (Figure 3H), consistent with being in the pachytene stage of meiosis (Figure 4A and Figure S4B).

**Figure 4.**
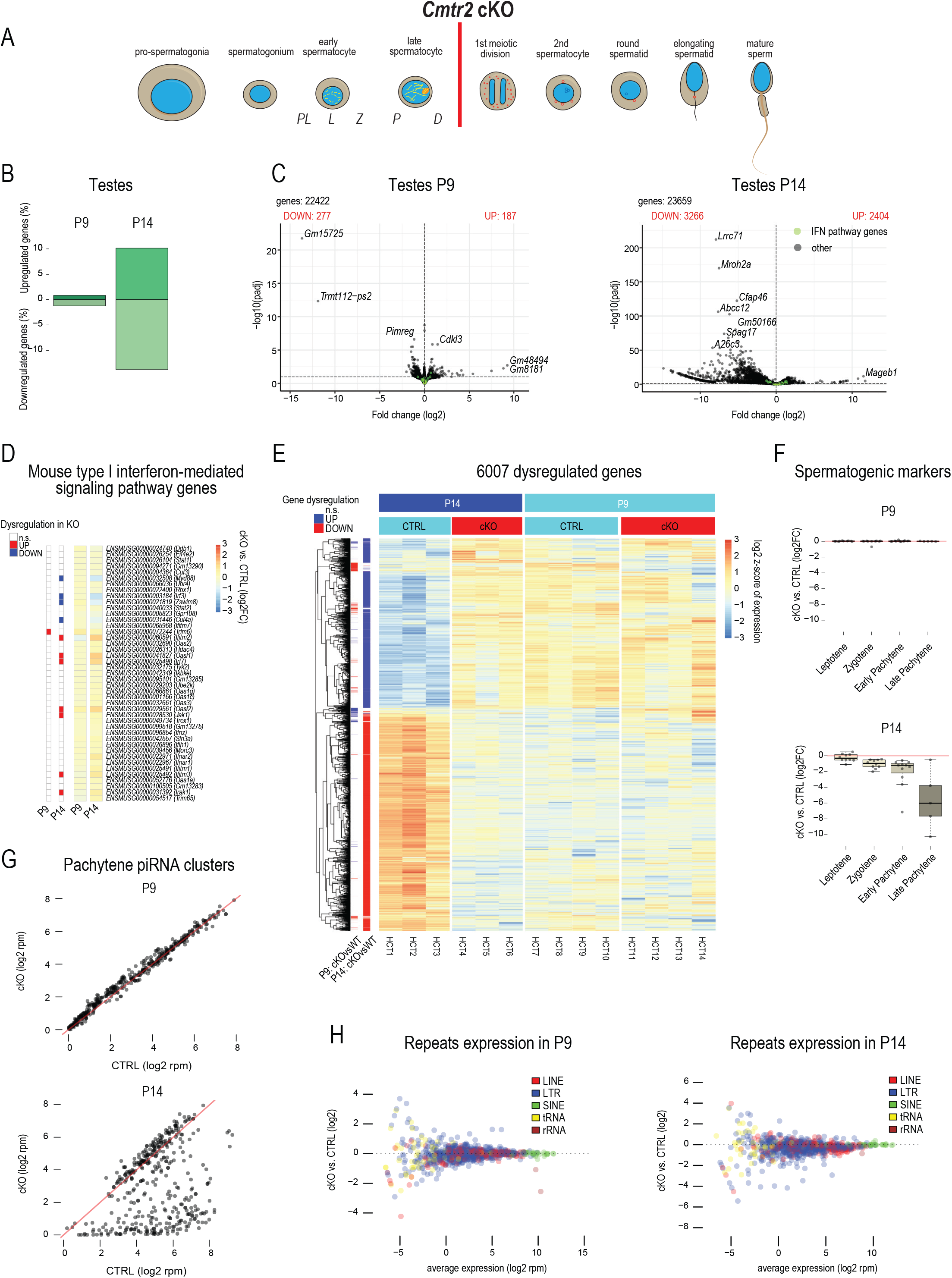
Cmtr2 knockout leads to arrest in mice spermatogenesis. (A) Schematic for spermatogenesis with arrest in *Cmtr2* knockout indicated by a red line. (B) Plot for the percentage of upregulated and downregulated genes in P9 and 14 *Cmtr2* cKO 4N spermatocytes (*Cmtr2^-/loxP^; DDX4-Cre^+/KI^*) as compared to the respective Control 4N spermatocytes (*Cmtr2^+/loxP^; DDX4-Cre^+/KI^*). (C) Volcano plot showing differential gene expression profile of the *Cmtr2* cKO spermatocytes compared to the Control spermatocytes from P9 (left) and P14 (right) mice. The genes associated with Interferon pathway are highlighted in green. (D) Heatmap showing the expression changes of Mouse type I interferon-mediated signalling pathway genes in *Cmtr2* cKO spermatocytes as compared to the respective Control spermatocytes (right). Differentially expressed genes (adjusted p-value < 0.1) are highlighted in red (upregulated) or blue (downregulated) (left). (E) Heatmap showing the expression pattern of all the dysregulated genes in Control and *Cmtr2* cKO spermatocytes from P9 and P14 mice (right). Differentially expressed genes (adjusted p-value < 0.1) are highlighted in red (upregulated) or blue (downregulated) (left). (F) Expression differences of meiosis I stage-specific marker genes (Leptotene, Zygotene, Early pachytene, Late pachytene) in *Cmtr2* cKO spermatocytes as compared to the respective control spermatocytes from P9 (top) and P14 (bottom) mice. (G) Correlation between the expression of individual pachytene piRNA clusters in *Cmtr2* cKO spermatocytes as compared to the respective control spermatocytes from P9 (top) and P14 (bottom) mice. (H) The MA plots displaying the expression changes of individual genomic repeats in *Cmtr2* cKO spermatocytes as compared to the respective control spermatocytes from P9 (left) and P14 (right) mice.

Transcriptome analysis shows that P9 control and *Cmtr2* cKO germ cells have largely similar gene expression profile with ∼2% of genes being dysregulated (Figure 4B). This contrasts with ∼25% of genes being altered in their expression levels in the P14 cKO germ cells (Figure 4B). Thousands of genes are altered in the P14 mutant germ cells, most being downregulated (Figure 4C and Figure S4C), with GO term analysis linking them to progression of spermatogenesis (Figure S4E). A heatmap representation of the gene expression changes between control and Cmtr2 cKO germ cells shows that gene expression at P9 stage leptotene-zygotene spermatocytes is largely similar (Figure 4E). Strikingly, mutant germ cells at P14, which based on molecular markers appear to be at pachytene stage (Figure 3G), have a transcriptome that closely resembles that in leptotene-zygotene spermatocytes (Figure 4E) resulting from failure to activate the P14 transcription program (Figure S4D). We collected a set of spermatogenic marker genes representative of different meiotic stages (leptotene to late pachytene) (Ernst *et al*, 2019) and examined their expression the P9 and P14 *Cmtr2* cKO germ cells (Figure 4F). Expression of such representative genes (STAR Methods) is similar in the control and *Cmtr2* cKO datasets in the P9 germ cells. Interestingly, expression of genes representative of leptotene spermatocytes is similar in the P14 control and *Cmtr2* cKO, but those representative of later stages (zygotene, early and late pachytene) are downregulated in the *Cmtr2* cKO (Figure 4F). This reinforces the conclusion that while germ cells in the *Cmtr2* cKO proceed to pachytene spermatocytes stage at P14 as determined by γ-H2AX and SYCP3 staining, their transcriptome fails to transition, resembling that of the preceding leptotene/zygotene stage.

We examined expression of genes involved in the Type I interferon-mediated signalling pathway, but did not observe any dramatic upregulation, with several genes being up- or down-regulated in P14 (Figure 4D). This is consistent with our observation in human KO cell lines and mutant liver and lung tissues (Figure 3). A recent related study (Zheng *et al*, 2026) examined *Cmtr2* cKO germ cells and proposed *Mov10l1* mRNA as a target for methylation-independent stabilization by CMTR2. MOV10L1 is a 5′-3′ RNA helicase that is essential for biogenesis of germline small RNAs called Piwi-interacting RNAs (piRNAs) that suppress transposable elements (Zheng *et al*, 2010). Mouse mutant for *Mov10l1* display male-specific sterility and de-repress transposable elements, none of which were observed in the *Cmtr2* cKO mice (Figure 4H and S4F). Instead, many genomic loci encoding for such small RNAs (pachytene piRNAs)(Li *et al*, 2013) that begin to be expressed in pachytene spermatocytes are mostly downregulated in the P14 dataset consistent with decreased expression of *Mybl1* transcription factor at P14 (Figure 4G and S4G). These data suggest a scenario where failure to activate the transcription program in P14 pachytene spermatocytes, which includes upregulation of many piRNA biogenesis factors like *Mov10l1* and the transcription factor *Mybl1* (Figure S4G) is the basis for the meiotic progression arrest. Taken together, we conclude that CMTR2 is required for meiotic progression by facilitating transcriptome transition that is required during germ cell development.

### *Cmtr2* is essential for definition of sex in zebrafish

Given that invertebrate Cmtr2 is biochemically (Haussmann *et al*., 2022) and genetically (Clemens *et al*., 2026; Haussmann *et al*., 2022) redundant with the Cap1 methylase, we wished to probe this question further with the zebrafish model. We used CRISPR to generate a 2.3 kb deletion within the *Dano rerio* (zebrafish) *Cmtr2* gene (STAR Methods and Figure S5A). Heterozygous *cmtr2^+/Δ^* fish were viable and fertile. Inter-crosses between the heterozygous parents gave rise to progeny that contained homozygous *cmtr2 ^Δ/Δ^* individuals (Figure 5A). Unlike the embryonic lethality in mice (Dohnalkova *et al*., 2023), *cmtr2* homozygous mutant zebrafish were viable and recovered in Mendelian ratios at 5 days post-fertilization (dpf) or as adults at 3.5 months (Figure 5A). Surprisingly, a majority of the heterozygous and most of the homozygous *cmtr2 ^Δ/Δ^* animals (hereafter referred to as the *cmtr2* mutant) were phenotypic males (Figure 5B). Homozygous *cmtr2 ^Δ/Δ^* animals were rarely identified as females. Histological examination of the testes from the control wildtype males reveal the characteristic pattern of seminiferous lobules which are outlined by a pale, eosinophilic layer of somatic interstitial cells that drapes around the germ cells within. The central cavity is full of spermatozoa that appear as basophilic granules. A similar observation is made for the homozygous animals, identifying them as males (Figure 5B). Such *cmtr2* mutant males were fertile, as they were able to produce progeny when crossed with wildtype or heterozygous *cmtr2^+/Δ^* females. We note that while many zebrafish germline factors when mutated result in a male-only phenotype, they also result in infertile individuals, as we demonstrated for *ythdc2* (Li *et al*, 2022). Thus, zebrafish Cmtr2 doesn’t seem to have a conserved role in meiotic progression, but has a role in definition of sex.

**Figure 5.**
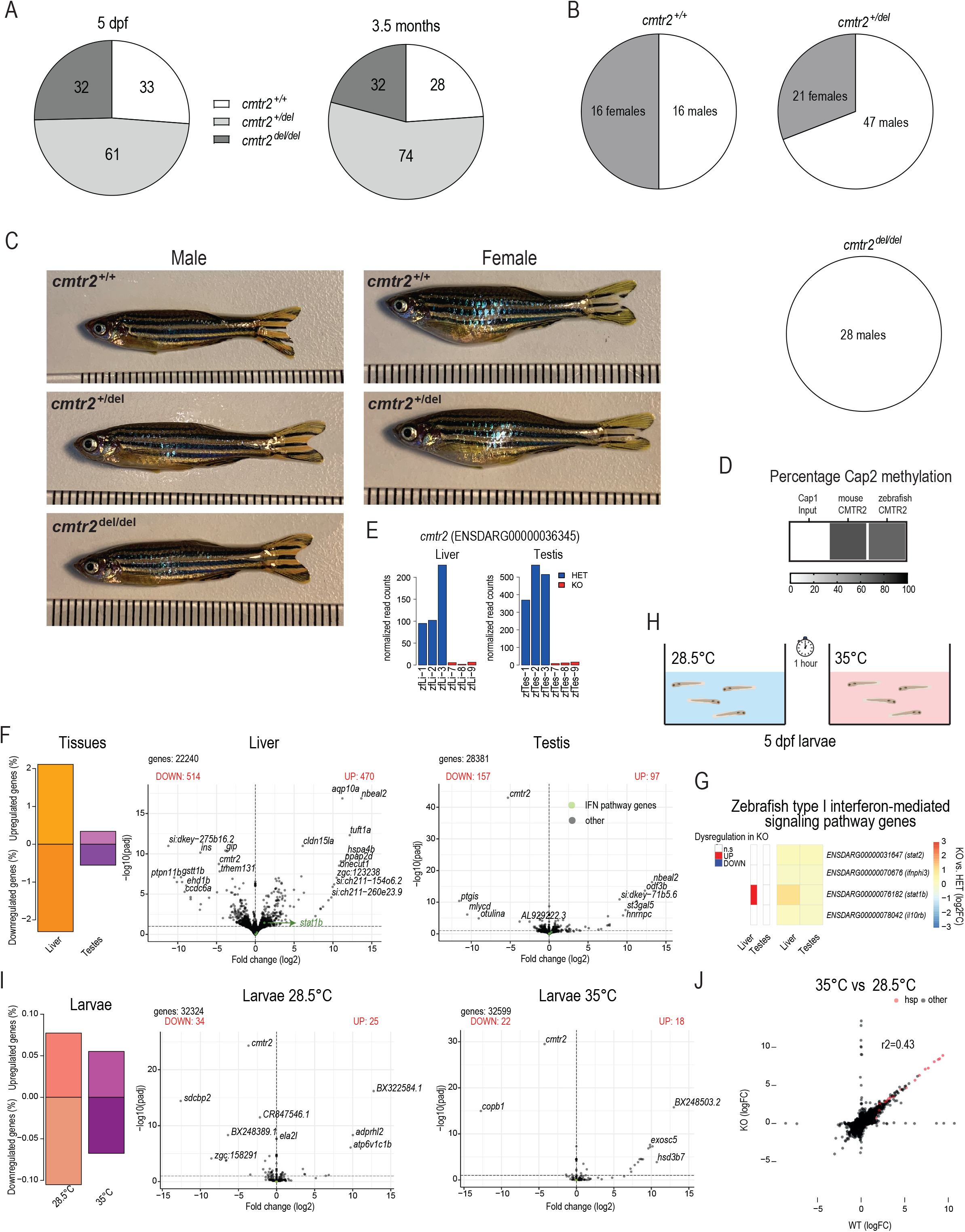
Cmtr2 knockout influences sex determination in Zebrafish. (A) Mendelian ratios in 5 days post fertilization (dpf) larvae and 3.5 month-old adult fish from heterozygous crosses. The larvae came from a mix of different crosses, the adults were a single cross. Individual fish numbers are shown. (B) Sex distribution in adult fish from a single heterozygote cross. Individual fish numbers are shown. Note: these are the same animals described in the right panel of A. 6 heterozygous fish were not sexed, hence there are 68 described here, and not 74. (C) Adult fish (8.5 months old) are phenotypically normal. Scale bar is a ruler with mm units. Males on the left, females on the right. (D) Recombinant mouse and zebrafish CMTR2 was used to *in vitro* methylate Cap1 RNAs substrate. After in vitro methylation, the RNA was purified and assessed by LC-MS to detect nucleotide 2’-O-methylation. Percentage Cap2 methylation was calculated upon normalization to the Cap1 methylated nucleotide. (E) Confirmation of *Cmtr2* knockout at RNA level. Expression levels of *Cmtr2* RNA in Control and *Cmtr2* KO zebrafish tissues. (F) Plot for the percentage of upregulated and downregulated genes in zebrafish *Cmtr2* KO tissues compared to the respective WT tissues (left). (Right) Volcano plots showing differential gene expression profile of the zebrafish *Cmtr2* cKO Liver/Testis as compared to the Control tissues. The genes associated with Interferon pathway are highlighted in green. (G) Heatmap showing the expression changes of Zebrafish type I interferon-mediated signalling pathway genes in zebrafish *cmtr2* cKO tissues as compared to the respective Control tissues (right). Differentially expressed genes (adjusted p-value < 0.1) are highlighted in red (upregulated) or blue (downregulated) (left). (H) Schematic for heat shock treatment of the zebrafish larvae 5dpf. (I) Plot for the percentage of upregulated and downregulated genes in zebrafish *cmtr2* KO larvae as compared to the respective WT tissues at different temperatures (left). (Right) Volcano plots showing differential gene expression profile of the zebrafish *cmtr2* cKO larvae as compared to the Control larvae at 28.5°C or 35°C. The genes associated with Interferon pathway are highlighted in green. (J) Correlation between gene expression changes in cKO larvae at 28.5°C and 35°C. The heat-shock genes are highlighted in red.

To formally confirm the biochemical activity of zebrafish Cmtr2, we prepared the recombinant protein (Figure S1B) and incubated with m7G-capped RNAs (STAR Methods). RNA mass spectrometry confirms that similar to the mouse CMTR2, the zebrafish orthologue prefers to ribose methylate the second nucleotide of m7G-capped RNAs (Figure 5D). Thus, zebrafish CMTR2 is a Cap2 methylase.

We isolated testes and liver from control and homozygous *cmtr2* mutant males and examined gene expression by RNAseq analysis. Only a small number of genes are dysregulated in the mutant testes (Figure 5F and Figure S5C). GO term analysis revealed enrichment of terms associated with the translation machinery among downregulated genes in the testes (Figure S5E). Given that the mutant animals are normally viable and fertile, it is unclear how these changes contribute to any physiological consequences. Mutant liver has much higher alteration of gene expression with ∼4% of genes dysregulated (Figure 5F and Figure S5D), with only DNA metabolic process (GO:0006259) term enriched among downregulated genes (Figure S5E). We examined the levels of four zebrafish genes annotated as Type I interferon-stimulated pathway genes, but these were not upregulated in the *cmtr2* mutant liver and testes (Figure 5G).

To examine whether zebrafish *cmtr2* has a role in stress response, we exposed 5 days post-fertilization (dpf) larvae to a shock increase in temperature from 28.5°C to 35°C for a duration of 1 hour (Figure 5H). Briefly, *cmtr2* mutant males were crossed with heterozygous *cmtr2^+/Δ^* females and a clutch of 24 larvae were exposed to the temperature shift, while a control set was left at 28.5°C. Some of the larvae did not survive the high temperature, but genotyping by RT-PCR did not show any preferential mortality of the *cmtr2* mutants. Transcriptome analysis indicates a clear and pronounced induction of heat shock proteins in wildtype and mutant larvae exposed to the temperature shock (Figure 5J). Overall, very few genes are altered in the mutants (Figure 5I and Figure S5G), with nuclear and nucleolar gene expression pathway being upregulated (Figure S5I), but this did not include the interferon pathway genes (Figure S5J). Taken together, we show that zebrafish *cmtr2* doesn’t have a conserved role in meiotic progression, similar to the mammalian orthologue, but is essential for definition of the female sex in zebrafish and has other gene regulatory roles under normal and stress conditions.

## DISCUSSION

The m7G cap is a signature feature of eukaryotic RNA pol II transcripts and this is critical for mRNA stability and translation. Cap1 methylation catalyzed by CMTR1 (Belanger *et al*., 2010) is a conserved feature in higher eukaryotes and serves as a discriminator between self vs non-self RNAs by preventing recognition by innate immune sensors (Inesta-Vaquera & Cowling, 2017). Cap2 mediated by CMTR2 (Werner *et al*., 2011) is also found in several organisms ranging from worms (Clemens *et al*., 2026) to flies (Haussmann *et al*., 2022) and humans (Despic & Jaffrey, 2023). A third modification, m^6^Am, is catalyzed by PCIF1 (Akichika *et al*., 2019; Boulias *et al*, 2019; Pandey *et al*, 2020; Sendinc *et al*, 2019), when the first nucleotide of the capped RNA is an adenosine.

In human cells, Cap1 and m6Am is almost ubiquitous on polyA+ RNAs (Akichika *et al*., 2019; Despic & Jaffrey, 2023), likely explained by the fact that nuclear CMTR1 and PCIF1 both carry WW domains that interact with the RNA pol II CTD, allowing them to gain constitutive access to the nascent RNA. Cap2 on the other hand is much more limited in polyA+ RNAs (Despic & Jaffrey, 2023; Dix *et al*., 2022). Here we show that CMTR2 is mostly nucleo-cytoplasmic (Figure 1D). Using an in vivo Cap2 sensor, we reveal that endogenous CMTR2 methylation activity is very limited and can be boosted by exogenously supplied CMTR2 (Figure 1C). Western analysis of endogenous CMTR2 from a knockin mouse line confirms that CMTR2 is a low abundant protein (Figure 1F). We speculate that CMTR2 levels are likely kept low in vivo.

A role for Cap2 in preventing activation of the innate immune pathway is proposed (Despic & Jaffrey, 2023). However, our own analysis with three independently generated CMTR2 KO human cell lines, three mouse KO tissues (Figure 2), and zebrafish KO tissues (Figure 5), does not support such a role for Cap2. Independently created CMTR2 KO A549 cells did not reveal innate immune activation (Aggarwal *et al*, 2026; Nukaga *et al*, 2025). It is possible that different genetic compensation mechanisms during establishment of mutant cell lines can be behind the discrepancy. Finally, a recent independently analysed germline cKO of mouse *Cmtr2*, did not reveal innate immune pathway activation (Zheng *et al*., 2026). Nevertheless, one study reported innate immune activation after treatment of cancer cells with a CMTR2-binder (Aggarwal *et al*., 2026). The authors used a modified DNA damaging agent (AT-1965), which was shown to bind CMTR2, but it is unclear if the agent inhibits RNA methylation activity of the enzyme.

Mouse *Cmtr2* is essential for embryonic development (Dohnalkova *et al*., 2023; Yermalovich *et al*., 2024), without any implication of the innate immune pathway. Our germline deletion of Cmtr2 reveals a role for the protein in transcriptome regulation that is required for meiotic progression (Figure 4). Similar findings were recently reported (Zheng *et al*., 2026). We show that CMTR2 is required for the germline transcriptome to transition, as the germ cells proceed along different stages of meiosis. The *Cmtr2* mutant germ cells fail to transition from the transcriptome as present in leptotene/zygotene spermatocytes, although the cells exhibit markers consistent with a successful passage into the pachytene stage (Figure 3).

Literature reveals how *Cmtr2* is used in different organisms for gene regulation. Invertebrates use it redundantly with *Cmtr1* (Clemens *et al*., 2026; Haussmann *et al*., 2022), but vertebrates (mice and zebrafish) seem to have dedicated roles for the two enzymes (Dohnalkova *et al*., 2023; Yermalovich *et al*., 2024). Although zebrafish *Cmtr2* is not involved in meiotic progression (at least in the male germline), it is critical for ensuring definition of the female sex, as all mutant animals are males (Figure 5). How CMTR2 or the Cap2 modification can influence gene expression is still an open question. RNA stability and translation was not shown to be affected in human cell lines (Despic & Jaffrey, 2023), but translation may be affected depending on the type of cell line used (Drazkowska *et al*., 2022). Cap2 is proposed to stabilize target RNAs in the mouse germline (Zheng *et al*., 2026) and our own work supports a role in transcriptome transition during germ cell development. Even splicing is proposed as a regulatory step for CMTR2 (Nukaga *et al*., 2025). Given the implication of CMTR2 in human cancers (Aggarwal *et al*., 2026; Cai *et al*, 2021; Nukaga *et al*., 2025), resolving this question of molecular role of Cap2 methylation for gene expression is a priority for future research.

## RESOURCE AVAILABILITY

### Lead Contact

Further information and requests for resources and reagents should be directed to and will be fulfilled by the Lead Contact Ramesh S. Pillai.

### Materials Availability

All unique reagents generated in this study, including the tagged-CMTR2 knock-in mouse line, are available from the Lead Contact without any restriction.

### Data and Code Availability

- Deep sequencing data generated in this study are deposited with Gene Expression Omnibus (GEO) under accession no. GEO: xxxxxxx.
- Code used in the current study is available from the Lead Contact upon reasonable request.

## ACKNOWLEDGEMENTS

We thank Nicolas Roggli for scientific illustration. We acknowledge the following University of Geneva core facilities: iGE3 Genomics Core; Transgenic Mouse Facility; Histology Facility. We also thank the EMBL Genomics core facility for deep sequencing; the Functional Genomics Center, Zurich for proteomics analyses. We thank the Karolinska Genome Engineering Facility, Karolinska Institutet for creation of the A549 CMTR2 KO cells. Carmen Fernandez Rodriguez was supported by a postdoctoral fellowship from the Lalor Foundation. This work was supported by grants to R.S.P. from Novartis Foundation for Medical-Biological Research (#24B138), the Swiss National Science Foundation (#310030_215346 and #310030_207468) and funding from the NCCR RNA & Disease (#51NF40_205601). Work in the Pillai lab is supported by the Republic and Canton of Geneva.

## AUTHOR CONTRIBUTIONS

H.R performed most of the experiments with mice and cell lines. E.D did initial in vitro methylation assays and characterized HEK293F CMTR2 KO cells; M.D initiated crosses for the Cmtr2 Cre-deletion lines; R.F created and characterized the fish mutant; C.B.V conducted RNA mass spectrometry; F.B helped with mouse experiments; C.F-R purified recombinant proteins; D.H conducted all computational analysis; R.S.P supervised the study and wrote the manuscript with input from everyone.

## STAR METHODS

### KEY RESOURCES TABLE

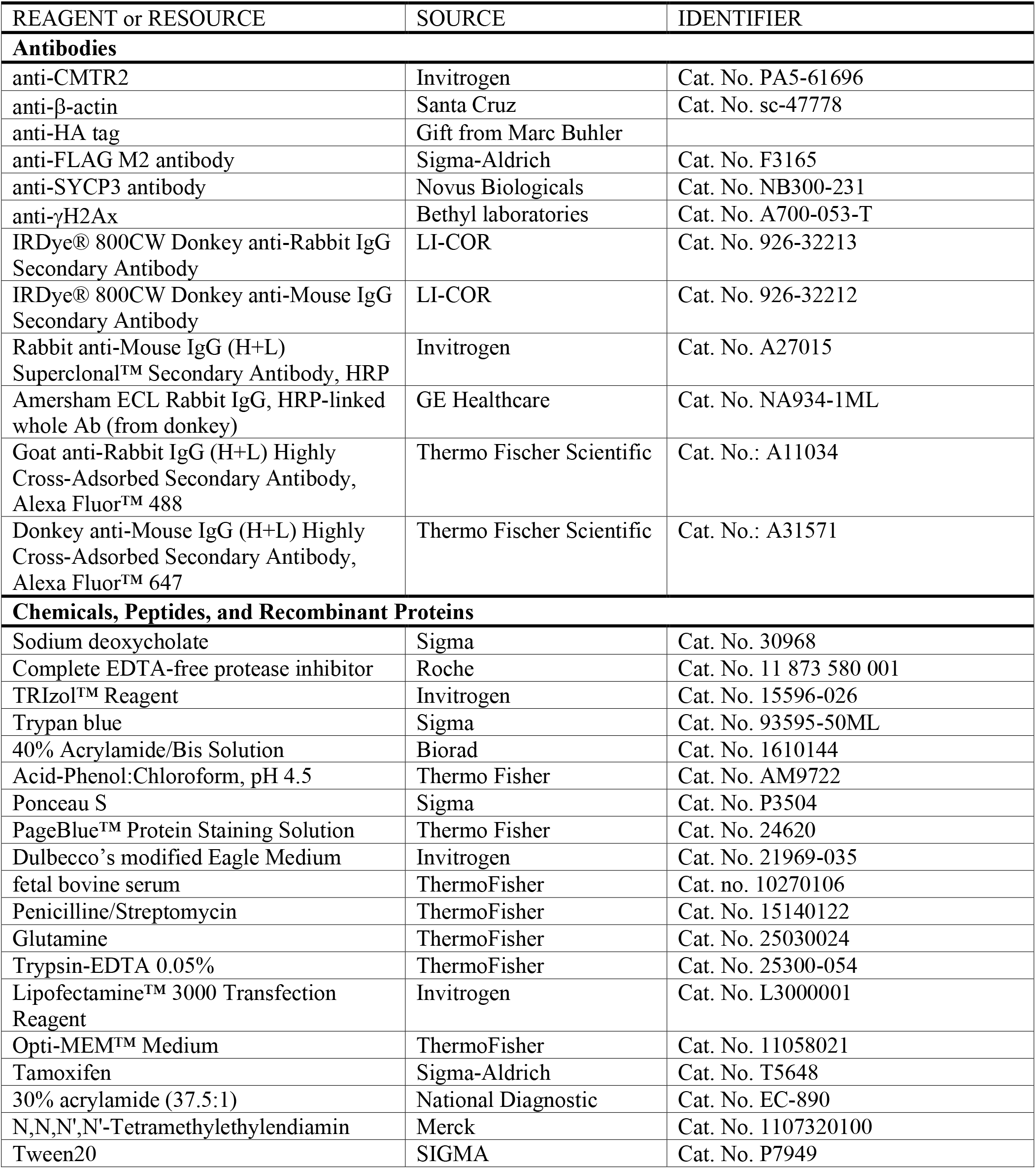
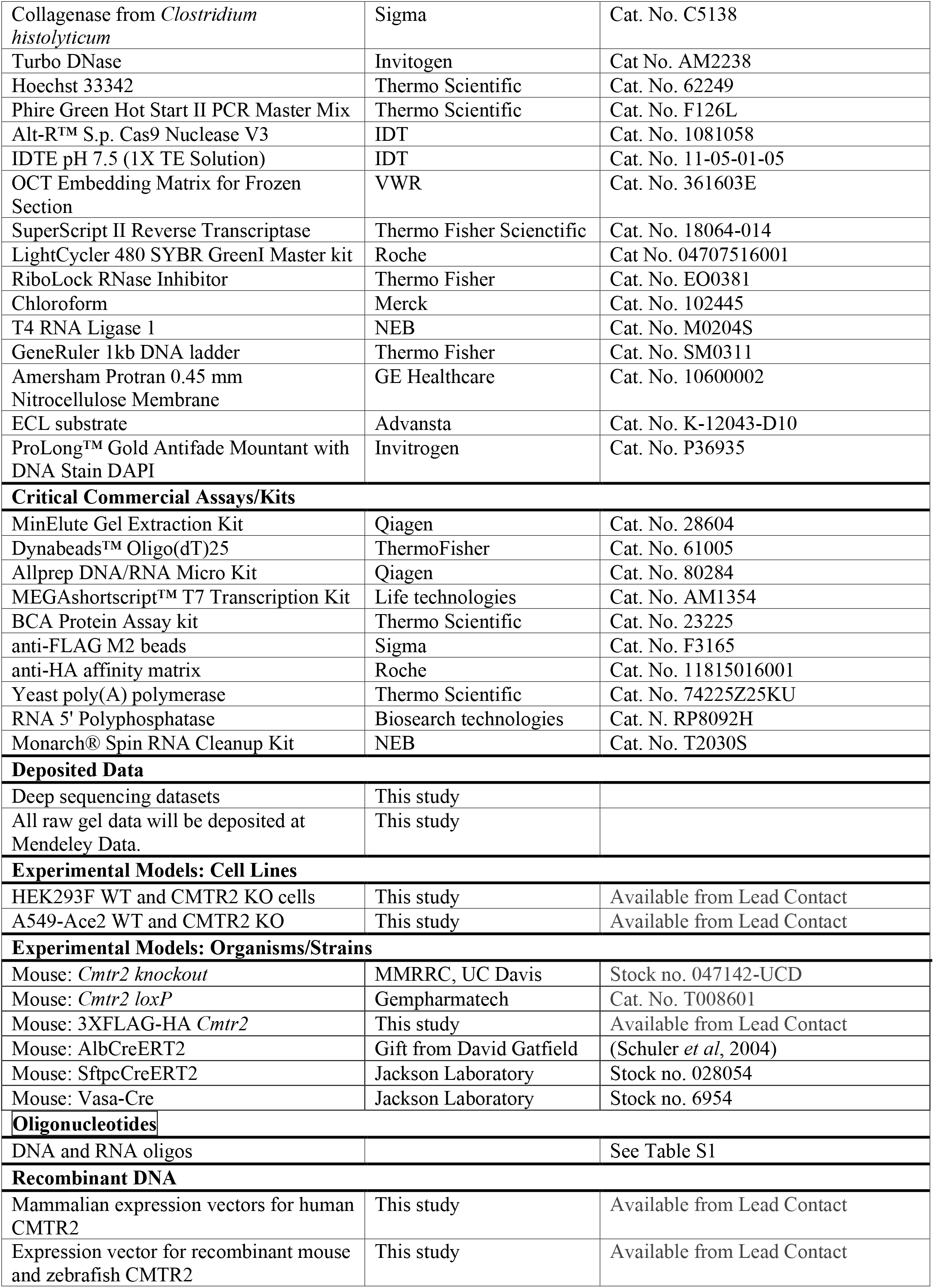
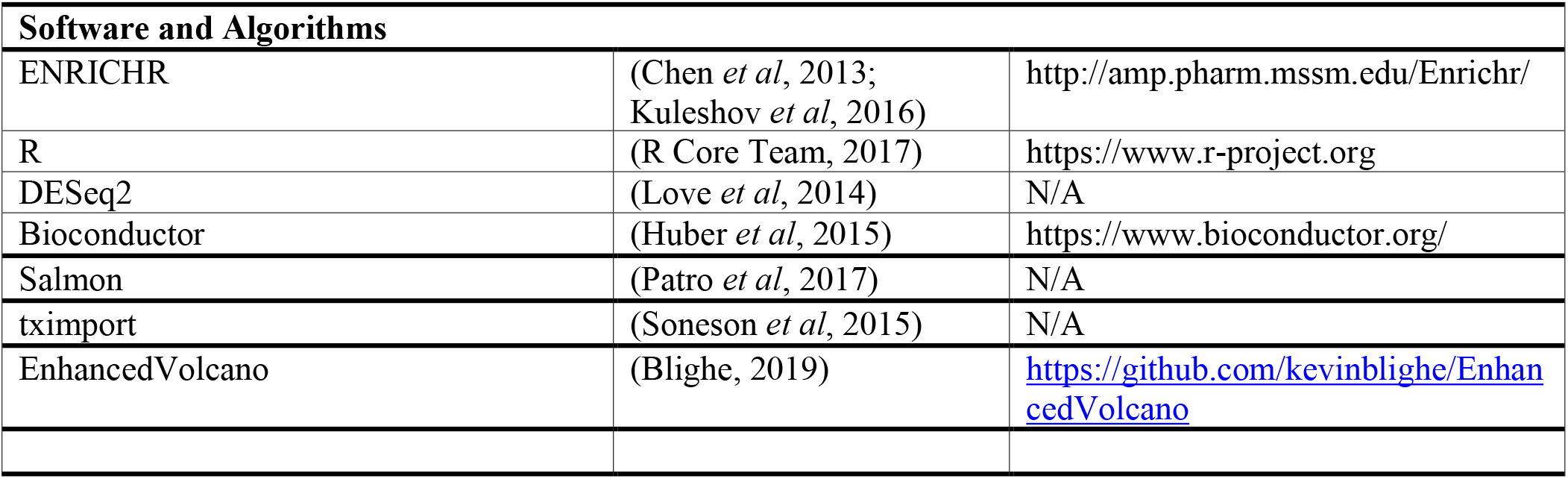

## EXPERIMENTAL MODEL AND SUBJECT DETAILS

The *Cmtr2* knockout model was generated by Knockout Mouse Project (KOMP) and the *Cmtr2* loxP floxed allele was generated by Gempharmatech, China. The N-terminal 3XFLAG-HA tagged Cmtr2 knock-in mouse line was generated at the Transgenic Mouse Facility at University of Geneva. Mice were bred in the Animal Facility of Sciences III at University of Geneva.

The use of animals in research at the University of Geneva is regulated by the Animal Welfare Federal Law (LPA 2005), the Animal Welfare Ordinance (OPAn 2008) and the Animal Experimentation Ordinance (OEXA 2010). The Swiss legislation respects the Directive 2010/63/EU of the European Union. Any project involving animals has to be approved by the Direction Générale de la Santé and the official ethics committee of the Canton of Geneva, performing a harm-benefit analysis of the project. Animals are treated with respect based on the 3Rs principle in the animal care facility of the University of Geneva. We use the lowest number of animals needed to conduct our specific research project. Discomfort, distress, pain and injury are limited to what is indispensable and anesthesia and analgesia is provided when necessary. Daily care and maintenance are ensured by fully trained and certified staff. This work was approved by the Canton of Geneva (GE294E).

### Mouse mutants

#### Cmtr2 knockout mouse

The *Cmtr2* knockout (KO) mutant mouse (C57BL/6N-*Cmtr2^tm1.1(KOMP)Vlcg^*/JMmucd, RRID: MMRRC_047142-UCD, Stock no. 047142-UCD) was generated as described (Dohnalkova *et al*., 2023).

### Cmtr2 loxP mouse

The Cmtr2 floxed mouse line was purchased from Gempharmatech (Catalogue no. T008601). The *Cmtr2* gene is located on mouse chromosome 8 and consists of two exons. The entire coding DNA sequence (CDS) is contained in Exon2. To create the conditional knockout mouse of *Cmtr2*, loxP sites were inserted in the same direction, flanking Exon2 (Figure S3A). Recombination of this region by Cre recombinase would result in the deletion of the intervening Exon2. The project was carried out by The Taconic-Cyagen Model Generation Alliance. Briefly, loxP site insertion was performed using CRISPR/Cas9 mediated genome engineering. Following two gRNAs were used for targeting the 5’ and 3’ region of Exon2:

gRNA 5S4: TCATGTACTGCCTCACCATA-GGG PAM

gRNA 3S4: TCCCCAGACAGCGCCACTAC-TGG PAM

Cas9, gRNA and targeting vector containing homologous arms and loxP sites flanking Exon2 in same direction were co-injected into fertilized eggs and the resulting pups genotyped for loxP sites insertion.

The *Cmtr2* loxP floxed mice of all genotypes and sexes were normally viable and fertile.

### 3X FLAG-HA KI mouse line

To detect the endogenous CMTR2 in mouse tissues, we tagged CMTR2 at the N-terminus with a 3X FLAG-HA tag. This mouse line was generated at the Transgenic Mouse Facility in University of Geneva. The *Cmtr2* gene is located on mouse chromosome 8 and consists of two exons. Exon 2 harbors the entire coding domain sequence (CDS) and was targeted in single cell mouse embryos from B6D2F1/J hybrid line (hybrid: 50% C57bl6; 50% DBA) using a specific guide RNA and single-stranded homology repair templates to insert the coding sequences for the 3X FLAG-HA tag, in-frame downstream of the ATG codon, at the N-terminal of CMTR2 protein (Figure 1C).

#### Purchase and preparation of gRNA

For gRNA preparation, crRNA and tracrRNA were annealed. Both these RNAs were purchased from Integrated DNA Technologies (IDT). The tracrRNA (67 bases, Cat. No. 1072533, IDT) is the part of gRNA that binds to the Cas9 protein and crRNA is the targeting part which has a complementarity of 20 bases with the genomic DNA of interest and some additional sequence for base-pairing with the 3’ end of the tracrRNA. The crRNA-tracrRNA base pairing region (GUUUUAGAGCUAUGCU) is common of generation of any gRNA. gRNA was generated by annealing 200 pmol each of tracrRNA and crRNA in IDTE buffer, pH7.5 (IDT, Cat. No. 11-05-01-05) to a final volume of 10 µl. The gRNA mix was heated to 95°C for 5 mins and thereafter allowed to cool for 30 mins in the heating block to obtain the functional annealed gRNA. The annealed gRNA was stored at -20°C till final injection.

*crRNA (CMTR2_Nterm_3XFLAGHA):*

/AltR1/AGCGCAGAAAGCUUCCGGCU GUUUUAGAGCUAUGCU /AltR2 /

AltR: chemical modifications to enhance stability of the RNA.

*Homology-dependent repair (HDR) template for 3X FLAG-HA tag (HDR_Nterm_CMTR2):*

AAGGTGACCCTTGTAACCTTCCGTTTTGTCTTCTCAGATTTTTAACTCAGTTTGACA**ATG**ga ctacaaagaccatgacggtgattataaagatcatgacatcgattacaaggatgacgatgacaagggcggcagcggctacccatacgatgttccaga ttacgctggtggaggcgggtctAGCA<u>AGCGCAGAAAGCTTCCGGCT</u>CGCCAGCCAGCTTGCTTGGAGA CCTTCAGCCCAGAC

The sequence in small letters represents the sequence corresponding to the N-terminal tag being inserted. The capital letters represent homology arms, with the gRNA sequence underlined. The initiator ATG for CMTR2 is marked in bold. The repair template was designed with a synonymous mutation to remove the gRNA binding site on the edited locus.

#### Preparation of injection mix

The annealed gRNA (final concentration of 0.6 pmol/μl) was mixed with the Cas9 protein (30 ng/μl, IDT, Cat. No. 1081058) in a final volume of 9 μl. The mix was incubated for 10 mins at room temperature for complex formation. The single-stranded HDR template (IDT, 20 ng/μl final concentration) was then added to the injection mix and the volume was adjusted to 100 μl with IDTE (IDT, Cat. No., 11-05-01-05). The injection mix was centrifuged for 5 mins at 13,000 rpm at 4°C and 50 μl of supernatant collected and stored on ice to be used for the injection.

Mouse embryos of hybrid background B6D2F1/J (black coat color) were injected with the mix at the Transgenic Mouse Core Facility, University Medical Centre (CMU), University of Geneva. The NMRI (Naval Medical Research Institute) mice (white coat color) were used as foster mothers.

The founder mice of hybrid background were identified by genotyping and backcrossed with wildtype C57Bl6/J (Janvier Labs) partners to obtain germline transmission. After two rounds of backcrossing, the heterozygous (*Cmtr2^+/KI^*) knockin (KI) males and females were intercrossed to obtain homozygous (*Cmtr2^KI/KI^*) knockin males and females. The homozygous males and females were further interbred to expand the colony, and the knockin males were used for CMTR2 pull down studies (Figure 1G and Figure S1F). This mouse line was designated as RP087: B6-*Cmtr2^em(3xFLAG-HA)Rspi^*/Ugfm.

The 3XFLAG-HA tagged *Cmtr2* knock-in mice of all genotypes and sexes were normally viable and fertile.

### Tamoxifen inducible Cmtr2 deletion in mouse liver/lungs

The Alb-CreERT2 mouse (Schuler *et al*., 2004) was used to generate liver specific tamoxifen-inducible *Cmtr2* knockout. This mouse specifically expresses tamoxifen-dependent Cre-ER^T2^ recombinase in hepatocytes (80% of total liver volume). This is achieved by inserting an IRES-containing coding region of Cre-ER^T2^ in the 3’ UTR of serum albumin gene. Alb-CreERT2 mouse was a kind gift from David Gatfield at University of Lausanne, Switzerland. Heterozygous *Cmtr2^-/+^* mice were crossed with *Alb-CreERT2^KI/KI^*mice to obtain *Cmtr2^-/+^; Alb-CreERT2^+/KI^* mice. These mice were crossed together to obtain *Cmtr2^-/+^; Alb-CreERT2^KI/KI^*which were finally crossed with *Cmtr2^loxP/loxP^* to obtain control (*Cmtr2^-/loxP^; Alb-CreERT^+/KI^*) and liver cKO (*Cmtr2^-/loxP^; Alb-CreERT2^+/KI^)* animals (n=3) upon intraperitoneal tamoxifen injection. Control and cKO females were used for the experiment.

The Sftpc-CreERT2 (Jackson Labs; Strain #028054) knock-in mouse was used to generate lungs specific tamoxifen inducible *Cmtr2* knockout. This mouse specifically expresses tamoxifen-dependent Cre-ER^T2^ recombinase in type II alveolar epithelial cells of lungs (60% of pulmonary alveolar epithelium). This is achieved by inserting an IRES-containing coding region of Cre-ER^T2^ in the 3’ UTR of the pulmonary-associated Surfactant protein C, *Sftpc* gene. Heterozygous *Cmtr2^-/+^* mice were crossed with *Sftpc-CreERT2^KI/KI^*mice to obtain *Cmtr2^-/+^; Sftpc^+/KI^* mice. These mice were crossed together to obtain *Cmtr2^-/+^; Sftpc-CreERT2^KI/KI^*which were finally crossed with *Cmtr2^loxP/loxP^* to obtain control (*Cmtr2^-/loxP^; Sftpc-CreERT2^+/KI^*) and lung cKO (*Cmtr2^-/loxP^; Sftpc-CreERT2^+/KI^)* animals upon intraperitoneal tamoxifen injection. Control and cKO males were used for the experiment.

20 mg/ml Tamoxifen (Sigma, Cat. No. T5648) was dissolved in corn oil (Sigma, Cat. No. C8267) overnight at room temperature, protected from light, on a cyclomixer. It was stored for upto 2 days at 4°C. Adult mice (P70) were injected with tamoxifen daily for five days and the mice were sacrificed after 6 days of last injection (i.e., 11 days after start of the experiment). Animals were euthanized by pentobarbital injection followed by cervical dislocation. Thereafter, livers and lungs were collected for experiments.

### Conditional deletion of Cmtr2 in the mouse germline

The *Ddx4-Cre (Vasa-Cre)* transgenic line (Stock No. 6954, Jackson Laboratory) was used generate germline specific *Cmtr2* knockout. This mouse specifically expresses Cre recombinase from germ cell-specific *Ddx4 (Vasa)* promotor (Gallardo *et al*, 2007). Multiple copies of the transgene are present in this line. Expression of the Cre recombinase begins at embryonic day 14.5 (E14.5) in both male and female germline. The animals obtained were back-crossed twice with wildtype C57BL/6J prior to establishing the *Cmtr2* mouse lines. *Cmtr2^-/+^; Ddx4-CreERT2^+/KI^* males were crossed with *Cmtr2^loxP/loxP^*females to prevent the maternal inheritance of Cre. The resulting Control (*Cmtr2^+/loxP^; Ddx4-CreERT2^+/KI^*) and cKO (*Cmtr2^-/loxP^; Ddx4-CreERT2^+/KI^*) animals were sacrificed at different time points to collect the testes (P9, P14, P75) and ovaries (P>60) for histological/transcriptomic analysis.

### Mouse genotyping

Ear punches were collected from weaned animals (at P21), and toe clips were collected for P4-P7 pups for genotyping. Earpunch/toeclips were digested in 100 μl of Lysis buffer (10 mM NaOH, 0.1 mM EDTA) at 95°C for 90 mins. The samples were centrifuged for 10 mins at 3000 rcf and 50 μl of supernatant transferred to a new tube containing 50 μl of TE buffer (20 mM Tris-HCl pH8.0, 0.1 mM EDTA). 2 μl of this digestion mix was used as template for PCR amplification. For the PCR reaction, 10 μl of Phire Green Hot Start II PCR Master Mix (Thermo Scientific, Cat. No. F126L), 2 μl of digestion mix and 1 ul each of the 10 mM Primer were made up to a volume of 20 μl with water. Specific PCR reactions were performed for different amplifications:

For amplification of *Cmtr2* Knockout allele, the primers MD280, MD282 and MD283 (Table S1) were used in the PCR reaction. Following conditions were used for PCR: 98°C for 30 s, 32 cycles of [98°C for 5 s, 65°C for 5 s and 72°C for 12 s], 72°C for 1 min, and finally at 4°C to hold the reaction. WT (244 bp) and KO (339 bp) PCR products were resolved on 2% agarose gel (Figure S3).

For amplification of *Cmtr2* loxP allele 5’ end, the primers MD408 and MD409 were used in a touchdown PCR reaction. Following conditions were used for PCR: 98°C for 30 s, 20 cycles of [98°C for 5 s, 65°C for 5 s (with 0.5°C decrease in each cycle) and 72°C for 12 s,], 20 cycles of [98°C for 5 s, 55°C for 5 s and 72°C for 12 s], 72°C for 1 min, and finally at 4°C to hold the reaction. WT (338 bp) and loxP KI (443 bp) PCR products were resolved on 2% agarose gel (Figure S3).

For amplification of *Cmtr2* loxP allele 3’ end, the primers MD410 and MD411 were used in a touchdown PCR reaction. Following conditions were used for PCR: 98°C for 30 s, 20 cycles of [98°C for 5 s, 65°C for 5 s (with 0.5°C decrease in each cycle) and 72°C for 12 s,], 20 cycles of [98°C for 5 s, 55°C for 5 s and 72°C for 12 s], 72°C for 1 min, and finally at 4°C to hold the reaction. WT (343 bp) and loxP KI (449 bp) PCR products were resolved on 2% agarose gel (Figure S3).

For amplification of *Sftpc-CreERT2* allele, the primers MD414, MD415 and MD416 were used in a touchdown PCR reaction. Following conditions were used for PCR: 98°C for 30 s, 10 cycles of [98°C for 5 s, 65°C for 5 s (with 0.5°C decrease in each cycle) and 72°C for 12 s,], 28 cycles of [98°C for 5 s, 60°C for 5 s and 72°C for 12 s], 72°C for 1 min, and finally at 4°C to hold the reaction. WT (343 bp) and loxP KI (449 bp) PCR products were resolved on 2% agarose gel (Figure S3).

For amplification of *Alb-CreERT2* allele, the primers MD366 and MD365 to amplify WT allele, and primers MD366 and MD367 to amplify the knockin allele were used in the PCR reaction. Following conditions were used for separate PCR reactions: 98°C for 30 s, 325cycles of [98°C for 5 s, 55°C for 5 s and 72°C for 12 s], 72°C for 1 min, and finally at 4°C to hold the reaction. Both reactions were pooled, and WT (229 bp) and KI (444 bp) PCR products were resolved on 2% agarose gel (Figure S3).

For amplification of *DDX4-Cre* allele, the primers MM113 and MM114 were used in the PCR reaction to amplify the transgene. Since insertion site of the transgene is not known, an internal control (IC) gene was amplified by the primers MD418 and MD419 in the same reaction. Following conditions were used for PCR: 98°C for 30 s, 35 cycles of [98°C for 5 s, 65°C for 5 s and 72°C for 12 s], 72°C for 1 min, and finally at 4°C to hold the reaction. WT (324 bp) and IC (349 bp) PCR products were resolved on 2% agarose gel (Figure S2). Since *DDX4-Cre* is a transgene, we can only confirm the presence of the transgene but cannot define its homozygous/heterozygous state by genotyping.

For amplification of *Cmtr2 N-term 3XFLAG-HA* insertion, the primers HR25 and HR26 were used in the PCR reaction. Following conditions were used for PCR: 98°C for 30 s, 32 cycles of [98°C for 5 s, 65°C for 5 s and 72°C for 12 s], 72°C for 1 min, and finally at 4°C to hold the reaction. WT (377 bp) and KO (497 bp) PCR products were resolved on 2% agarose gel (Figure S3).

The sequence information of all the primers is available in Table S1 and the representative gels are shown (Figure S1D and Figure S3B-D).

### Zebrafish Cmtr2 KO

A mutated *cmtr2* allele with a 2.3 kb deletion was generated by targeting *cmtr2* with CRISPR-Cas9 using two sgRNAs as shown (Figure S5A). sgRNAs were designed to target sequences in the *cmtr2* gene, at the 5’ end of the coding region with a sequence complementary to 5’-GTGATGAGGAGATTCGTGCA-3’ and within the 3’UTR with the equivalent RNA sequence to 5’-TAGTGCCACGGTGGACATTT-3’. Ribonucleoprotein complexes of each were prepared with recombinant Cas9 nuclease, mixed, and microinjected into early embryos of the TU genetic background, essentially as described previously (Li *et al*., 2022). Fish were raised and crossed to identify founders, bred to produce heterozygotes, and then families raised to assess allele distribution. Fish were genotyped using PCR reactions on fin-clips using allele discriminatory oligonucleotides: cmtr2_genF 5’-GATGTCACCAAACTTGAGTTCAAAG-3’ and cmtr2_genR 5’-AATCCCAGTCATATGCAGCACATTC-3’.

### CMTR2 KO Cell lines

#### A549-ACE2 CMTR2 KO

We used A549 adenocarcinomic human alveolar basal epithelial cells that express the human ACE2 receptor (A549 Ace2) cells, which were a kind gift from the laboratory of Prof. Volker Thiel, University of Bern. A549-Ace2 *Cmtr2* KO cell lines were generated at the Karolinska Genome Engineering Facility, Karolinska Institutet. Briefly, two synthetic sgRNAs were used for CRISPR-Cas9 mediated gene editing. The sgRNAs were precomplexed with Cas9 protein the complexed ribonucleoproteins (RNPs) were electroporated (Neon system, ThermoFisher Scientific) in A549-Ace2 WT cells. Guide RNAs were g2: 5’-GTGATAAGAAACTGGATGAG-3’ and g3: 5’-GAAATGCATTAAGTTCAGTG-3’

Cells were expanded for several days prior to genomic DNA extraction and used for estimating frequency of edited alleles in a dropoff assay (probes for the WT allele will dropoff if the site was edited) using droplet digital PCR. Cell pools with a 60% dropoff for the edited CMTR2 locus were expanded and single cells in 96-well plates were genotyped for the edited region. A549 cells are hypotriploid, as they contain three incomplete sets of chromosomes. Two KO clones (RP16 and RP39) which contained edits that are out of frame (small deletions and insertions) in all three alleles were used for further experiments, after confirmation with sanger sequencing. PCR primers used for amplification are provided (Table S1). The knockout status was also confirmed by western blotting (Figure S2A).

#### KO clone 1

RP-16. It contains a -9 bp deletion, which was the combination of a -10 bp deletion which led to a STOP codon, followed by a +1 insertion. The other two alleles were also both out-of-frame.

#### KO clone 2

RP-39. It contains 3 alleles with insertions (+1, +58, +76) and one allele with -17 bp deletion.

#### HEK293F CMTR2 KO

Human embryonic kidney cells HEK293F, which are suitable for suspension cultures, were used for editing the *CMTR2* locus (Synthego). One sgRNA was used for CRISPR-Cas9 mediated gene editing. The sgRNA was precomplexed with Cas9 protein the complexed RNPs were electroporated in HEK293F cells. sgRNA: 5’-AACUGCCACUCAUUAUUAAG-3’

Single cell dilution was performed from the edited pool obtained upon analysis of sequencing data from PCR amplified fragments by Synthego’s Interference of CRISPR Edits (ICE) software tool. The sorted cells were expanded and genotyped for the edited region. The final clone used as HEK293F CMTR2 KO was homozygous for a +1 insertion in CMTR2 coding region. A cell pool obtained from mock-transfected cells was used as HEK293F WT cell line. PCR primers used for amplification is provided (Table S1).

## METHOD DETAILS

### Plasmid constructs and transfection

#### Plasmids for mammalian cell expression

##### pCI-neo-3XFLAG-HA-hCMTR1

The codon optimized CDS for human CMTR1 coding region (NCBI: NP_055865.1) with N-terminal 3XFLAG-HA tag coding region was synthesized in pCI-neo vector. *pCI-neo-3XFLAG-HA-hCMTR2*: The codon optimized CDS for human CMTR2 coding region (NCBI: NP_001311307.1) with N-terminal 3XFLAG-HA tag coding region was synthesized in pCI-neo vector. *pCI-neo-hPCIF1-3XFLAG-HA*: The codon optimized CDS for human PCIF1 coding region (NCBI: NP_071387.1) with C-terminal 3XFLAG-HA tag coding region was synthesized in pCI-neo vector. *pcDNA3.1-GFP-hCMTR2*: The codon optimized CDS for human CMTR2 with N-terminal EGFP coding region was synthesized in pcDNA3.1 vector.

##### pcDNA3.1-NLS-GFP-hCMTR2

The coding sequence for nuclear localization sequence (NLS) from CMTR1 (aa residues 2-19) was inserted at the N-terminal of EGFP in pcDNA3.1-GFP-hCMTR2 construct.

##### pcDNA3.1-GFP-NLS-hCMTR2-WW

The coding sequence for WW domain from CMTR1 (aa residues 755-790) was inserted at the N-terminal of EGFP in pcDNA3.1-GFP-NLS-hCMTR2 construct.

Lipofectamine3000 (Invitrogen, Cat. No. L3000001) was used for the transfection of the constructs in mammalian cells as per manufacturer’s protocol.

### Plasmids for recombinant protein expression

Mouse CMTR2 coding sequence (aa 15-759, NCBI: NP_666327.2) was cloned into vector pACEBac2-His-Strep-SUMO, for protein production in insect cell expression system, as described previously (Dohnalkova *et al*., 2023). The full-length zebrafish CMTR2 coding sequence (NCBI Ref Seq. XP_073762474.1.) was cloned into the same vector for expression in insect cells.

### Purification of mouse and zebrafish CMTR2 protein

Mouse CMTR2 was purified as described (Dohnalkova *et al*., 2023).

To purify of zebrafish CMTR2, the pACEBac2-His-Strep-SUMO-zCMTR2 was transformed into DH10EMBacY competent cells for preparation of the bacmids. The bacmid DNA was extracted and transfected with FuGENE 4K (Promega, Cat. No. E5911) into the Sf9 insect cells for virus production. The supernatant (V0) containing the recombinant baculovirus was collected after 72 h post-transfection. To amplify the virus pool, 3 mL of the V0 virus stock was added into 25 mL of Sf9 (0.5 × 10^6^/mL) cells. The resulting cell culture supernatant (V1) was collected 24 h after cells entered proliferation arrest. For large-scale protein production, Hi5 cells were infected with virus (V1) and cells were harvested 72 h post-infection.

Hi5 cells expressing full-length zebrafish CMTR2 were resuspended by lysis buffer (40 mM imidazole, 50 mM Tris-HCl pH 8.0, 300 mM NaCl, 5% glycerol and 5 mM 2-mercaptoethanol) supplemented with proteinase inhibitor (Thermo Scientific, EDTA-free). Cells were disrupted by sonication and then cell lysate was centrifuged at 18,000 rpm for 50 min at 4°C. The clarified supernatant was incubated with Ni^2+^ chelating Sepharose HPbeads (Cytiva, Cat. No. 17526801) at 4°C for 2 h. After incubation, beads were washed with an imidazole gradient in the wash buffer (40 mM or 50 mM imidazole, 50 mM Tris-HCl pH 8.0, 300 mM NaCl, 5% glycerol and 5 mM 2-mercaptoethanol). Proteins bound to the beads were eluted with elution buffer (50 mM Tris-HCl pH 8.0, 300 mM NaCl, 250 mM Imidazole, 5% glycerol and 5 mM 2-mercaptoethanol).

The eluate was subjected to a second affinity purification step using Strep-Tactin column (IBA, Cat. No. 2-1201-010), with bound protein eluted using 5 mM desthiobiotin in buffer: 50 mM Tris-HCl pH 8.0, 300 mM NaCl, 5% glycerol and 5 mM 2-mercaptoethanol. The N-terminal 6xHis-Strep-SUMO-TEV tag was cleaved overnight with TEV in the dialysis buffer (25 mM Tris-HCl pH 8.0, 150 mM NaCl, 5% glycerol and 5 mM 2-mercaptoethanol). After dialysis, cleaved tag was removed via second nickel column and flow-through containing the untagged protein was pooled and concentrated using a Pierce Protein Concentrator (ThermoScientific, Cat. No. 88529). The concentrated protein was further purified by size-exclusion chromatography on a Superose 6 Increase 10/300 GL (GE Healthcare) pre-equilibrated with the buffer (25 mM Tris -HCl pH 8.0, 150 mM KCl, 5% glycerol, 2 mM DTT). Peak fractions were analyzed by SDS-PAGE electrophoresis (Figure X) and fractions containing pure protein were supplemented with 10% glycerol, flash-frozen in liquid nitrogen and stored at −80°C. Pure untagged mouse and zebrafish CMTR2 proteins were used for in vitro methylation assays (Figure S1A-B).

### Antibodies used in this study

#### Primary antibodies

Rabbit anti-CMTR2 (Invitrogen, Cat. No. PA5-61696); mouse anti-actin (Santa Cruz, Cat. No. sc-47778); rabbit anti-SYCP3 (Novus Biologicals, Cat. No. NB300-231); rabbit anti-γ-H2AX (Bethyl laboratories, Cat. No. A700-053-T); mouse monoclonal anti-HA (a kind gift of Marc Buehler, FMI, Basel); mouse anti-FLAG (Sigma-Aldrich, Cat. No. F3165).

#### Secondary antibodies

For Western blot analyses, the following secondary antibodies conjugated to Horse Radish Peroxidase were used: anti-rabbit IgG HRP-linked (GE Healthcare, Cat. No. NA934), anti-mouse IgG HRP-linked (Invitrogen, Cat. No. A27015). Fluorescent probe-conjugated secondary antibody used were IRDye® 800CW Donkey anti-Rabbit IgG Secondary Antibody (LI-COR, Cat. No. 926-32213) and IRDye® 800CW Donkey anti-Mouse IgG Secondary Antibody (LI-COR, Cat. No. 926-32212).

For immunofluorescence, anti-rabbit Alexa Fluor 488 antibody (ThermoFisher Scientific, Cat. No. A11034) and anti-mouse Alexa Fluor 647 antibody (ThermoFisher Scientific, Cat. No. A31571).

Anti-HA affinity matrix (Roche, Cat. No. 11815016001) and anti-FLAG M2 beads (Sigma, Cat. No. F3165) were used for immunoprecipitations.

### Germ cell sorting from testes

Testes were collected from P9, P14 animals in PBS and decapsulated using the forceps. The tubules were transferred to prewarmed tubes containing 1 mg/ml type IV Collagenase (Sigma, Cat. No. C5138) prepared in DMEM, supplemented with DNase I (Invitrogen, Cat. No. AM2238) (10 μl DNAseI for 1.5 ml Collagenase). The tubes were incubated on the shaking incubator at 300 rpm for 15 mins at 35°C. After 5 mins of shaking, the tubules were gently pipetted for 2-4 times. After centrifugation at 278 RCF for 5 mins at room temperature, the pellet was resuspended in prewarmed 0.125% Trypsin (Gibco, Cat. No. 25200-056) containing 0.5 mg/ml Collagenase-DNAse I and placed on the shaking incubator at 300 rpm for another 7 mins at 35°C. The tubules were gently pipetted midway of incubation. After incubation, 1 ml of DMEM supplemented with 10% FBS was added to the tube and gently mixed. After centrifugation at 278 RCF for 5 mins at room temperature, the pellet was resuspended in FACS buffer (1% BSA-PBS) and filtered through 40 μm cell strainer (Merck, CLS352340). Cell viability was checked with trypan blue, which stain dead cells. Cells were incubated with DNA-staining dyes Hoechst 33342 (Thermo Scientific, Cat. No. 62249) and DRAQ7 (stains DNA in only dead or damaged cells) (Thermo Scientific, Cat. No. D15105) for sorting. Cell sorting of meiotic 4n spermatocyte population was performed using the cell population gates described previously (Gaysinskaya & Bortvin, 2015) using FACS BDAria at the Flow cytometry Core facility, University Medical Centre (CMU), University of Geneva.

### RNA extraction from sorted cells and sequencing

4N Spermatocytes were collected in FACS buffer and pelleted at 278 x g for 5 mins at 4° C. RNA was extracted from the pelleted cells using Allprep DNA/RNA micro kit (Qiagen, Cat. No. 80284). The extraction was performed according to the manufacturer’s protocol and elution was done in 10 μl DEPC-water.

RNA quantification was performed using Qubit 2.0 fluorometer (Invitrogen, SN 2286611551) and RNA integrity was assessed on TapeStation (Agilent). 10 ng of total RNA from sorted cells was used as input for library preparation. NEBNext UltraII Directional RNA library Prep Kit for Illumina (NEB, Cat. No. Cat. No. E7760L) was used for library preparation in combination with NEBNext rRNA Depletion kit V2 (NEB, Cat. No. E7400L). The amounts and quality of libraries were assessed using Tapestation (High sensitivity D100 ScreenTape, Agilent Technologies, Part No. 5067-5584). Equimolar libraries were pooled and sequenced on NovaSeq X Plus 1.5 billion flow-cell for 100 bp paired-end reads at the EMBL Genomics Core Facility.

### Marker genes for spermatogenesis stages

The gene list used as meiotic stage specific markers was curated from (PMID:30890697).

Leptotene: Prss50, Smc3, tex15, Smc1b, Syce2, Dmc1, Prdm9, Fbxw2, Brca2, Rec8 Zygotene: Ly6k, Sycp3, Sycp1, Tex101, Emc7, Rad51ap2, Syce3, H2afx, Sycp2, Meiob Early pachytene: Clgn, Hormad1, Cntd1, Tdrd9, Piwil2, ATR, Scmh1, Catsperb, Kdm3a, Msh4, Mybl1 Late pachytene: Spata16, Spink2, Pgam2, Nme5, Tex40

### RNA isolation from mouse tissues and library preparation for sequencing

For total RNA extraction from mouse lungs and liver, frozen tissue (in liquid nitrogen) was minced to a fine powder using mortal and pestle. 30 mg minced tissue was used for RNA isolation and added to 1 ml of TRIzol reagent (ThermoFisher Scientific, Cat. No. 15596018) in a 1.5ml microcentrifuge tube. The tissue was homogenized in TRIzol reagent using micropestle (Carl ROTH, YE14.1). Chloroform (200 μl) was added and the tube and vortexed for 40 sec. The tubes were centrifuged at 12000 rpm for 20 mins at 4°C. The aqueous layer was collected and equal volume of isopropanol added. The solution was mixed gently, kept at room temperature for 5 mins and centrifuged at 12000 rpm for 30 mins at 4°C. The pellet was washed with 70% ethanol, air-dried and resuspended in nuclease-free water. DNase I (ThermoFisher Scienctific, Cat. No. AM2238) treatment was done for 30 mins at 37°C. The volume of the solution was made up to 200 ul with nuclease-free water and equal volume of acidic Phenol: Chloroform (1:1) was added. The tubes were vortexed for 40 sec and then centrifuged at 12000 rpm for 10 mins at 4°C. After the spin, aqueous layer was collected to a new tube where 1/10^th^ volume of 3M sodium acetate (pH5.2) and 2.5 times cold 100% ethanol was added, the solution mixed gently and incubated at -80°C overnight. The following day, RNA was precipitated by centrifugation at 12000 rpm for 30 mins at 4°C. The RNA pellet was washed with 70% ethanol, air-dried and resuspended in nuclease-free water.

1 μg of total RNA from liver and lung tissues was used as input for library preparation. NEBNext UltraII Directional RNA library Prep Kit for Illumina (NEB, Cat. No. E7760L) was used for library preparation in combination with NEBNext rRNA Depletion kit V2 (NEB, Cat. No. E7400L). The amounts and quality of libraries were assessed using Tapestation (DNA High sensitivity chip). Equimolar libraries were pooled and sequenced on NextSeq2000 P3 for 50 bp paired-end reads at the EMBL Genomics Core Facility.

### cDNA synthesis and Real time PCR

*Cmtr2* RNA expression in liver and lungs from Control and *Cmtr2* cKO mice was quantified using real time PCR. 2 μg of isolated RNA was used for cDNA synthesis using gene-specific reverse primers (Table S1) for *Cmtr2* and *β-actin* (internal control). cDNA synthesis was done using SuperScript II Reverse Transcriptase (Thermo Fisher Scienctific, Cat. No. 18064-014) as per manufacturer’s protocol.

*Cmtr2:* Annealing temp.: 58°C, Amplicon size: 299 bp. *β-actin:* Annealing temp.: 58°C, Amplicon size: 189 bp.

Quantitative RT-PCR (qRT-PCR) was done using LightCycler 480 SYBR GreenI Master kit (Roche, Cat. No. 04707516001). 2 μl of cDNA was used for the qRT-PCR in a 20 μl reaction according to manufacturer’s protocol for 45 cycles. The comparative threshold cycle (CT) method was used to calculate fold change in *Cmtr2* RNA levels (2^-ΔΔCT^) upon normalisation with β-actin (Figure 2I and 2K).

### Collection of mouse tissues for western blot

Whole tissue/cell lysates were prepared from tissues collected from mice. A piece of tissue was homogenized in 1X RIPA lysis buffer (50 mM Tris pH 7.4, 150 mM NaCl, 0.25% sodium deoxycholate, 1% NP-40, 1 mM EDTA, Complete Protease Inhibitor Cocktail tablet (Roche, Cat. No. 5056489001) using a dounce homogenizer. The suspension was incubated on a cyclomixer for 1 hr at 4°C, tubes centrifuged at 10,000 rpm for 10 mins and the supernatant collected.

### Western blot

Protein concentration was quantified by using BCA Protein Assay kit (Thermo Scientific, Cat. No. 23225). The samples were boiled for 10 min in SDS loading buffer and resolved via SDS-PAGE. Proteins from the gel were transferred onto a nitrocellulose membrane (Cytiva) using a semi-dry transfer apparatus (Biorad), overnight at 5V. After incubation with the desired primary antibodies, blots were incubated with either HRP-tagged or fluorescent secondary antibodies. For HRP-tagged secondary antibodies, the blots were developed using ECL substrate (Advansta, Cat. No. K-12043-D10) in a ChemiDoc and for fluorescent antibodies, the blots were dried after washing and scanned (784 nm laser combined with 829/62 nm bandpass filter module-784/829BP62) directly using Sapphire FL Biomolecular Imager (Azure Biosystems, Inc., S/N: SFL-1044).

### Immunoprecipitation of endogenous FLAG/HA-CMTR2 from mouse tissues

Tissues from *3XFLAG-HA Cmtr2* mouse line were used for immunoprecipitation (IP) either using magnetic anti-FLAG M2 beads (Sigma, Cat. No. F3165) or anti-HA affinity matrix (Roche, Cat. No. 11815016001). The lysate for IP was prepared by homogenizing the tissue in 1X lysis buffer [50 mM Tris pH8.0, 150 mM NaCl, 0.5 mM EDTA, 0.1% NP40, 10% glycerol, Complete Protease Inhibitor Cocktail tablet (Roche, Cat. No. 5056489001)], similar to the lysate preparation for western blot. The lysate was incubated with anti-FLAG M2 beads (for FLAG-IP) or anti-HA affinity matrix (for HA-IP) for 3 hrs at 4°C to allow the binding of protein to the beads. 50 μl of beads was used for each pull down reaction. After incubation, the beads were washed 5 times with lysis buffer (10 min incubation at 4°C on cyclomixer for each wash). From FLAG beads, the proteins were eluted using 3XFLAG peptide (0.25 mg/ml, Bachem, H-7536.0005) for 1hr at 4°C. From HA beads, the proteins were eluted by boiling the beads in electrophoresis buffer (20 mM Tris pH7.5, 2 mM EDTA, 5% SDS, 200 mM DTT, 20% glycerol, bromophenol blue). 20 μl of eluates were loaded on the 10% SDA-PAGE for western blot (Figure 1G, S1F).

### Histology, immunofluorescence staining of mouse tissues

For histological analysis, the dissected tissues were washed in 1X PBS and fixed in 4% paraformaldehyde at 4°C overnight. Next day, after washing with PBS, the tissues were transferred to the embedding cassette (Simport, Cat. No. M508-3) and paraffin embedded at the Histology Core Facility of CMU, University of Geneva. After paraffin embedding, 7 μm sections were cut and stained with hematoxylin and eosin. The sections were examined under the Zeiss Axio Imager M2 microscope and images taken.

For immunofluorescence staining of tissues, the tissue samples were embedded in Optimal cutting temperature compound (OCT) and 7 μm sections cut using cryostat and slides stored at -80°C. For immunofluorescence staining, the sections were air-dried, fixed with 4% PFA for 10 mins at room temperature. After washing with PBS, the sections were permeabilized using 0.1% Triton-X-100 in PBS for 10 mins at room temperature. The tissued were blocked in the blocking solution (10% goat serum, 1% BSA, 0.1M glycine in 1X PBS) and stained overnight with primary antibodies at 4°C. Next day, after washing with PBS, the slides were stained with anti-rabbit Alexa Fluor 488 antibody for 1 hr at room temperature. The slides were washed and mounted using ProLong Gold with DAPI (Thermo Fischer Scientific, Cat. No. P36931). The images were taken using the Zeiss LSM800 confocal microscope at the Bioimaging centre at University of Geneva.

### Imaging in transfected human cells

HEK293F cells were seeded on coverslips in a 24-well plate in DMEM media supplemented with 10% FBS. 14-16 hrs after seeding, the cells were transfected with the desired construct using Lipofectamine3000. 24h after transfection, the coverslips were washed with cold PBS and fixed with 4% PFA for 20 mins at room temperature. After washing with PBS, the coverslips were permeabilized using 0.1% Triton-X-100 for 5 mins at room temperature.

For cells expressing 3XFLAG-HA tagged constructs, the coverslips were blocked using 3% BSA-PBS and stained with anti-FLAG antibody overnight at 4°C. Next day, after washing with PBS, the anti-mouse Alexa Fluor 647 antibody for 1 hr at room temperature. The slides were washed and mounted using ProLong Gold with DAPI and imaged.

For cells expressing GFP-tagged constructs, the coverslips were mounted with ProLong Gold with DAPI after permeabilization. The images were taken using Zeiss LSM800 confocal microscope at the Bioimaging centre at University of Geneva.

### In vitro transcription of Cap2 sensor RNAs

We wished to prepare m7G-capped Cap2 sensor RNAs that carry a heavy isotope-labelled second nucleotide, which can then be specifically detected in RNA isolated from transfected cells. Methylation status of this second nucleotide (Cap2 methylation) can be quantified by RNA mass spectrometry. Two different m7G-capped and polyadenylated RNAs were prepared. Sensor I was without any open-reading frame, while Sensor II was a mRNA of Renilla luciferase.

### Sensor I sequence

m7GpppGA*UCUUCCUCUCUCUUCUCCUUCUCCUCUCUUCUCCCUUCUCUCC This sensor RNA is a noncoding RNA, designed to have a unique nucleotide at the second position, which is marked a with heavy isotope-labelled residue (A*).

The DNA oligos containing a T7 promoter sequence upstream of the desired RNA sequence were synthesized (Table S1) and annealed by heating at 95°C for 5 mins and allowing the reaction to cool to room temperature at the ramp speed of 0.1 °C/s in annealing buffer (10 mM Tris pH7.5, 100 mM NaCl).

2 μM of this template was used for setting up the *in vitro* transcription (IVT) reaction using MEGAshortscript kit (Invitrogen, Cat. No. AM1345) as per manufacturer’s protocol. The IVT reaction was supplemented with 75 mM m^7^G(5’)ppp(5’)G RNA Cap Structure Analog (NEB, Cat. No. S1404S) and 75 mM heavy isotope-labelled ATP (made from ^15^N, ^13^C instead of ^14^N and ^12^C) (Silantes, Cat. No. 121603611) to produce a Cap0 RNA containing heavy A (heavier by 12 Daltons, compared to the naturally occurring adenosine in cells) at the second nucleotide. The reaction was purified after DNase I digestion, using Monarch® Spin RNA Cleanup Kit (NEB, Cat. No. T2030S) and polyadenylated using Yeast poly(A) polymerase (Thermo Scientific, Cat. No. 74225Z25KU) to add a poly(A) tail of around 100 nts. After final purification, the RNA integrity and size was checked using 8% Urea-PAGE gel ***Sensor II:***

With the aim of obtaining a coding RNA with the sensor segment, the sensor I (prior to poly (A) tailing) was ligated to another *in vitro* transcribed RNA coding for luciferase (Figure 1C).

Luciferase RNA was *in vitro* transcribed using linearized phRL-Tk vector. Regular NTPs and no cap analog was used for this reaction, resulting in a 5’ triphosphate luciferase RNA, which was polyadenylated using poly(A) polymerase. This RNA was treated with RNA 5’ Polyphosphatase (Biosearch technologies, Cat. No. RP8092H) to obtain a 5’ monophosphate RNA for ligation.

Both the Sensor I RNA and the luciferase RNA were ligated using a splint DNA (Table S1) and T4 RNA ligase I (NEB, M0204S) for 2 hrs at 37°C. The sequence of the splint oligo was such that after hybridization with the two RNAs to be ligated, there will be a 6 nts RNA overhang at the 3’ end and 1 nt at the 5’ end. This has been shown to increase the ligation efficiency of RNAs (Kim *et al*, 2023). The ratio of 5’ RNA (capped Sensor I RNA with heavy A at second nucleotide): 3’ RNA (polyadenylated luciferase RNA): Splint = 2:1:2 was used for ligation. The RNA was purified after verification in a 4% Urea-PAGE gel (Figure x). The ligated RNA was generated with around 50% efficiency and was used for transfection experiments.

### PolyA+ RNA purification

After transfection of cells with the sensor RNAs, the cells were harvested at the desired time points and total RNA isolated using TRIzol reagent, as described previously. The total RNA was used to isolate polyA+ RNA which was analysed by RNA mass spectrometry. For the polyA+ RNA purification, Dynabeads™ Oligo (dT)25 (Invitrogen, Cat. No. 61002) were used as per manufacturer’s protocol and the RNA was eluted in 10 μl nuclease free water.

### In vitro methylation with recombinant CMTR2

The 20 nt RNAs for *in vitro* methylation reaction were prepared as described previously using MEGAshortscript kit (Invitrogen, AM1354). Sequence DNA templates used are provided (Table S1). 500 pmol capped RNAs were incubated with 0.5 µm recombinant mouse /zebrafish CMTR2 in a reaction containing 2.5 mM SAM and 40 units of RNase inhibitor (Thermo Scientific, EO0381) in the methylation buffer (25 mM Tris pH7.5, 50 mM KCl). The reaction was incubated at 37°C for 16 hrs and the methylated RNA purified using Monarch® Spin RNA Cleanup Kit (NEB, T2030L) and analysed by mass spectrometry to quantify the levels of methylation (Figure 1B, 5D).

### Quantifications of RNA modifications using LC-MS

RNA mass spectrometry was performed as described previously (Dohnalkova *et al*., 2023).

## QUANTIFICATION AND STATISTICAL ANALYSIS

Sequence analysis was as described previously (Dohnalkova *et al*., 2023).

## SUPPLEMENTAL INFORMATION

Supplemental Information includes 5 figures and can be found with this article online at.

---

**Figure S1:**
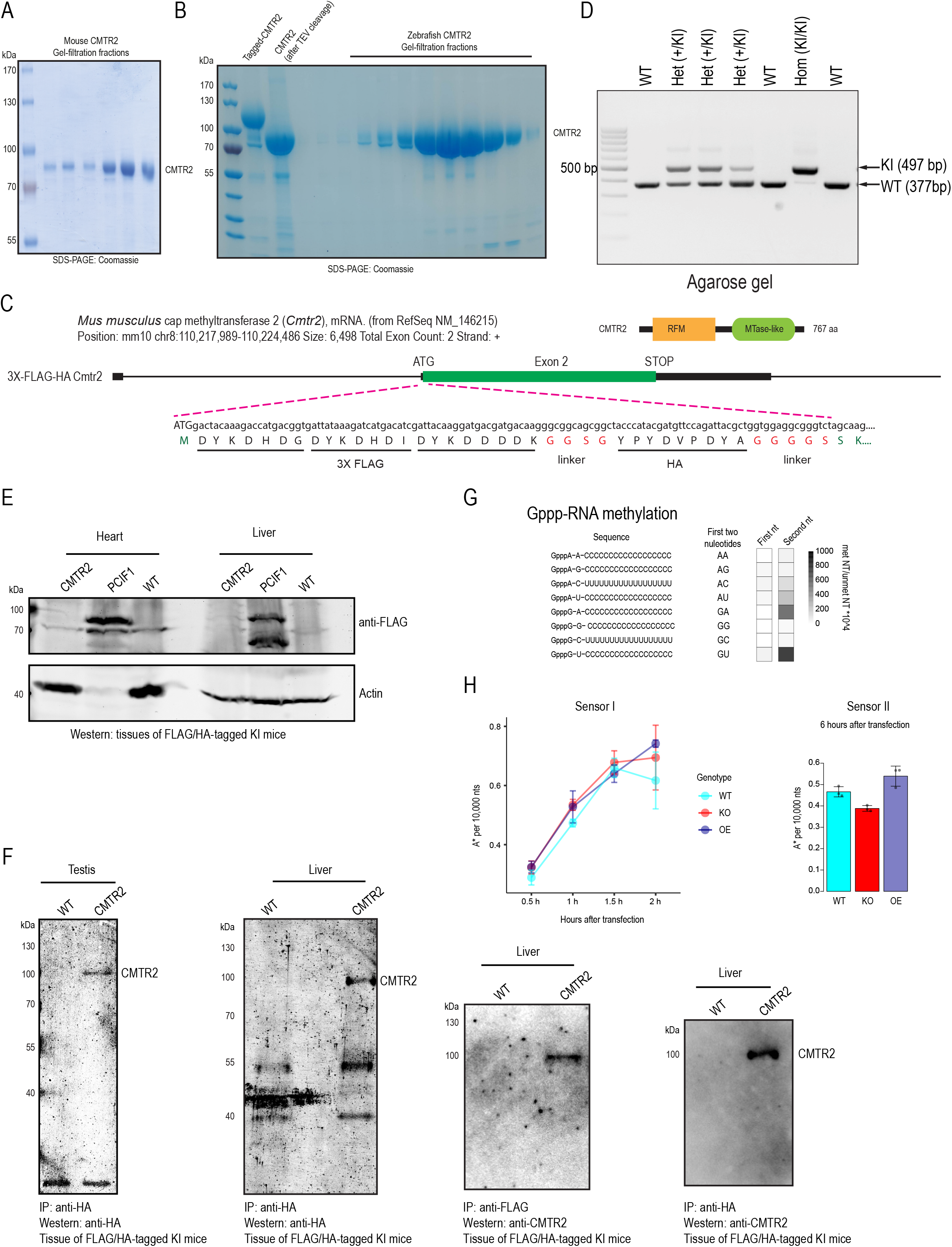
Generation of conditional *Cmtr2* knockout lines (*Cmtr2* cKO). (A) Coomassie stained gel for the purified mouse CMTR2 protein. (B) Coomassie stained gel for the purified zebrafish CMTR2 protein. (C) Schematic for the 3XFLAG-HA tag inserted at the N-terminus of CMTR2 coding region. (D) Agarose gel showing the genotyping PCR strategy used to identify the wildtype and tagged mouse. (E) Tissue expression analysis of *3XFLAG-HA-Cmtr2* (CMTR2) knock-in mouse. *Pcif1-3XFLAG* (PCIF1) mouse tissues were used as positive control and wild type (WT) mouse tissues were used as negative control. Western blotting was performed using anti-FLAG antibody. β-actin was used as the loading control. (F) Immunoprecipitation of CMTR2 was performed from Liver and Testis of the *3XFLAG-HA-Cmtr2* (CMTR2) knock-in mouse using anti-FLAG /anti-HA beads. Tissues from WT mouse were used as control. Western blotting was done using anti-CMTR2/ anti-HA antibodies as described. (G) *In vitro* methylation using mouse CMTR2 protein. The RNA substrates with the described sequences were used for in vitro methylation and the methylation of first and second nucleotide quantified by LC-MS. (H) The levels of sensor RNAs was determined by quantifying the abundance of heavy A nucleotide (A*) at the described time points.

**Figure S2:**
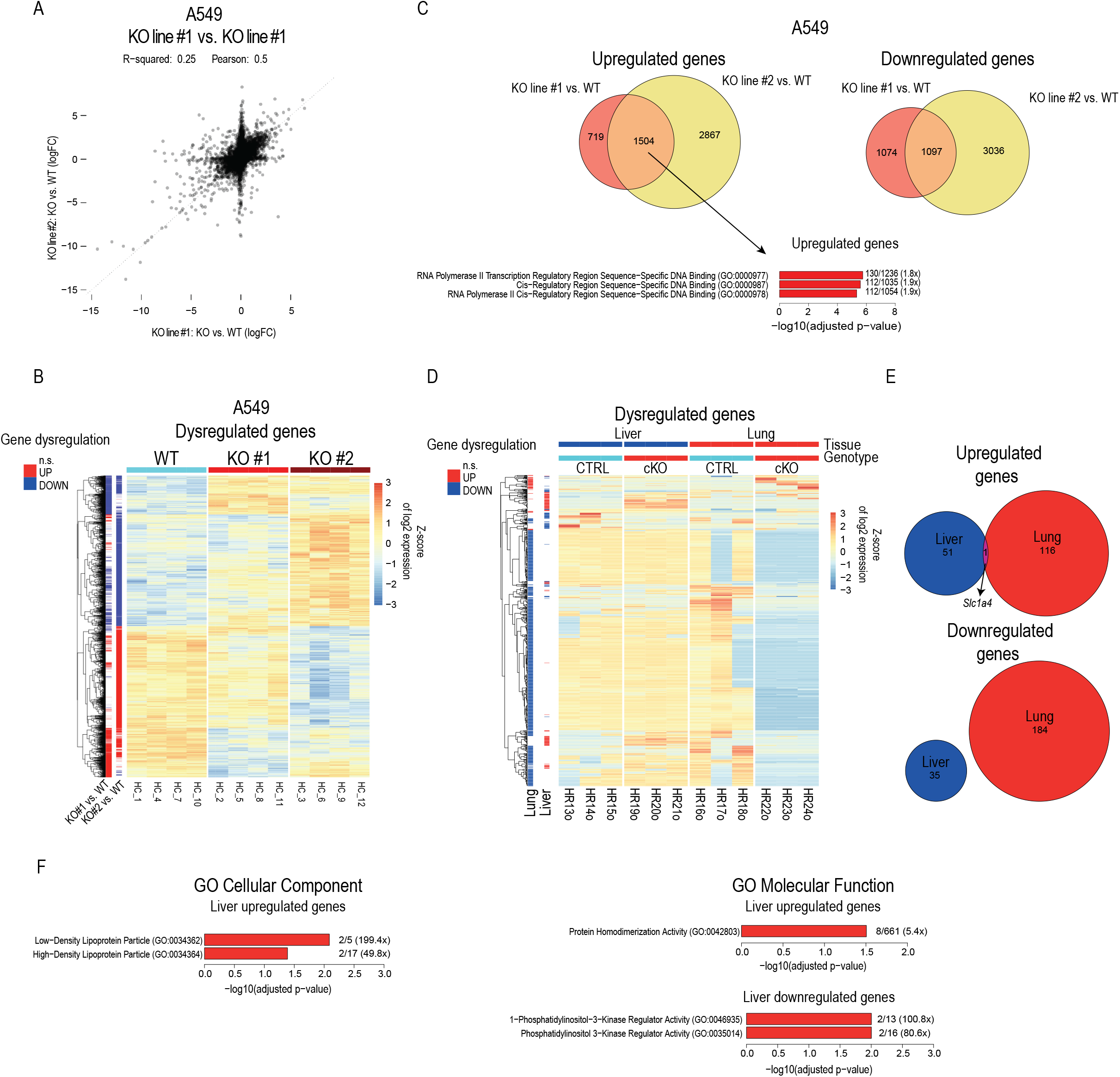
Gene expression alterations upon *Cmtr2* knockout. (A) Correlation between gene expression changes among two *Cmtr2* KO A549-Ace2 cell lines (KO line#1, KO line#2). (B) Heatmap showing the expression pattern of all the dysregulated genes in WT and *Cmtr2* KO A549-Ace2 cell lines (KO#1, KO#2) (right). Differentially expressed genes (adjusted p-value < 0.1) are highlighted in red (upregulated) or blue (downregulated) (left). (C) Venn diagrams were generated to determine the common upregulated and downregulated genes in the two A549-Ace2 *Cmtr2* knockout cell lines. The enriched gene ontology categories for the common upregulated genes are shown. (D) Heatmap showing the expression pattern of all the dysregulated genes in Control and *Cmtr2* cKO liver and lung tissues from adult mice (right). Differentially expressed genes (adjusted p-value < 0.1) are highlighted in red (upregulated) or blue (downregulated) (left). (E) Venn diagrams were generated to determine the common upregulated and downregulated genes in the mouse liver and lung samples upon *Cmtr2* cKO. (F) The enriched gene ontology categories for the upregulated and downregulated genes in liver are shown.

**Figure S3:**
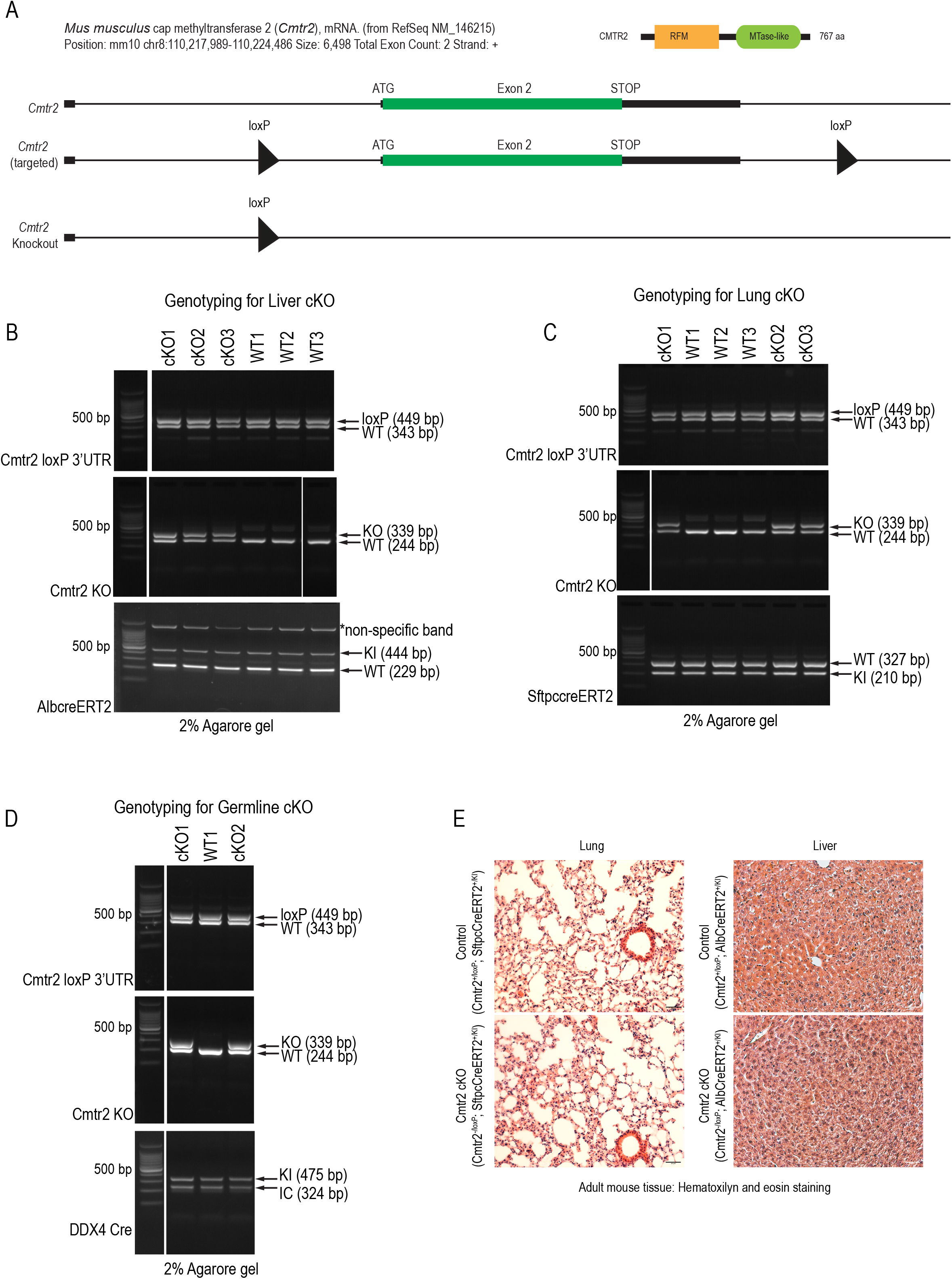
Characterization of *Cmtr2* cKO mouse lines. (A) Schematic for the generation of *Cmtr2* loxP mouse line. These mice were bred with the desired Cre lines to generate *Cmtr2* conditional knockouts (cKO). (B) Agarose gels showing genotyping PCR strategy used to identify liver-specific Control (*Cmtr2^+/loxP^; AlbCreERT2^+/KI^)* and *Cmtr2* cKO (*Cmtr2^-/loxP^; AlbCreERT2^+/KI^)* mice. (C) Agarose gels showing genotyping PCR strategy used to identify lung-specific Control (*Cmtr2^+/loxP^; S*ftpcCreERT2*^+/KI^)* and *Cmtr2* cKO (*Cmtr2^-/loxP^; S*ftpcCreERT2*^+/KI^)* mice. (D) Agarose gels showing genotyping PCR strategy used to identify germline-specific Control (*Cmtr2^+/loxP^; DDX4-Cre^+/KI^)* and *Cmtr2* cKO (*Cmtr2^-/loxP^; DDX4-Cre^+/KI^)* mice. (E) Histological analysis of hematoxylin and eosin-stained sections of adult Control and *Cmtr2* cKO Lung (left) and Liver (right). Scale bar represents 50 μm.

**Figure S4:**
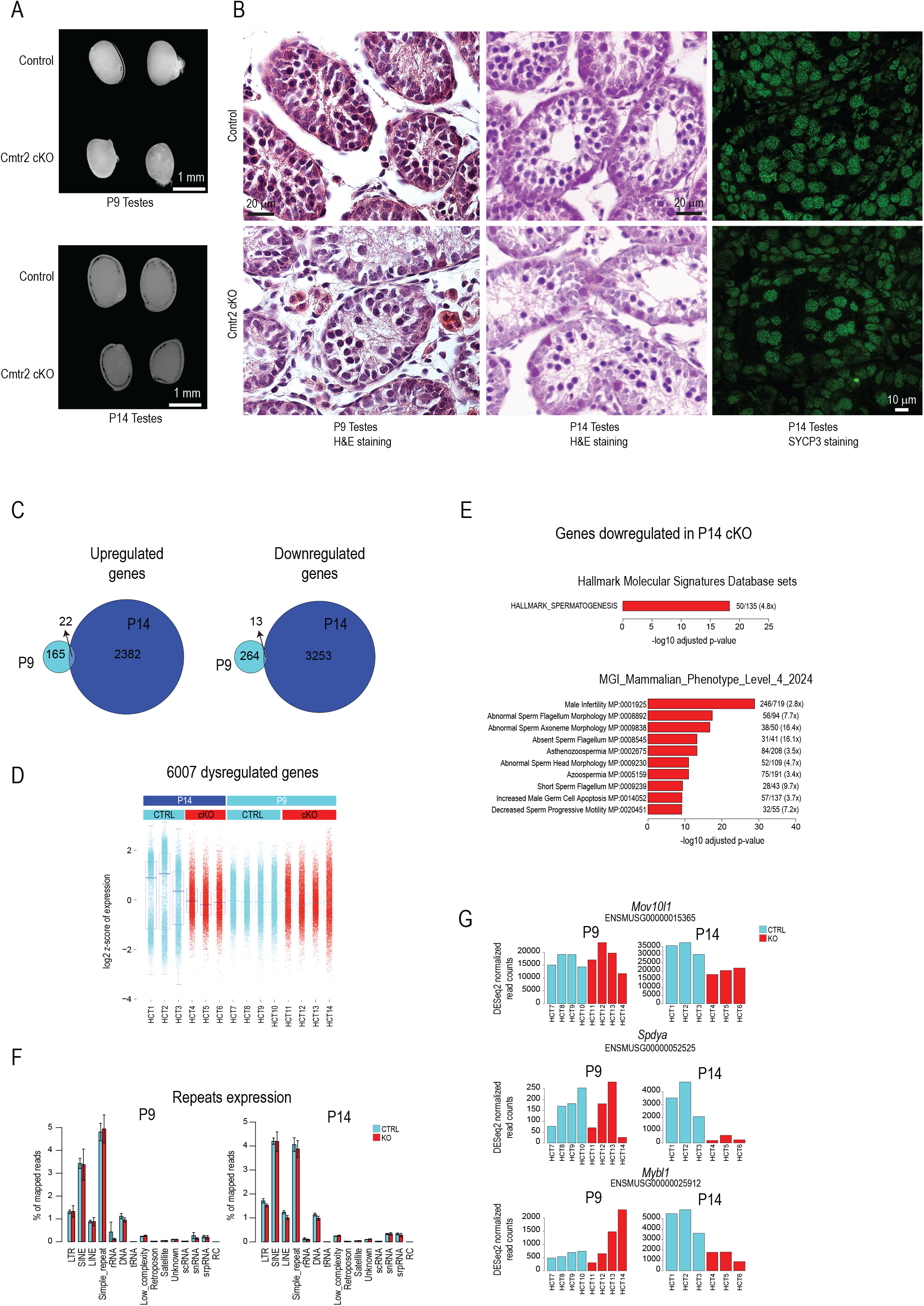
Characterization of gene expression alterations upon *Cmtr2* cKO in male germ cells. (A) Testes size upon *Cmtr2* cKO. Comparison of Control and *Cmtr2* cKO testes of animals aged P9 and P14. Scale bar represents 1 mm. (B) Histological analysis of hematoxylin and eosin-stained sections of Control and *Cmtr2* cKO testes from P9 and P14 animals. Scale bar represents 20 μm as indicated. SYCP3 staining in Control and *Cmtr2* KO testes (P14). The testicular sections from Control and *Cmtr2* cKO animals were used for immunofluorescence staining using anti-SYCP3 primary antibody and AF488-conjugated secondary antibody (Green). Scale bar represents 10 μm. (C) Venn diagrams were generated to determine the common upregulated and downregulated genes in the mouse spermatocytes upon *Cmtr2* cKO in P9 and P14 mice. (D) Boxplot comparing the expression of all dysregulated genes between *Cmtr2* cKO spermatocytes as compared to the respective control spermatocytes. (E) The gene ontology categories enriched in downregulated genes in P14 mouse spermatocytes upon *Cmtr2* cKO are shown. (F) Expression of individual repeat classes does not change upon *Cmtr2* cKO in spermatocytes from P9 and P14 mice. (G) Expression levels of specific genes (*Mov10L1, Spdya, Mybl1*) in the Control and *Cmtr2* cKO spermatocytes from P9 and P14 mice.

**Figure S5:**
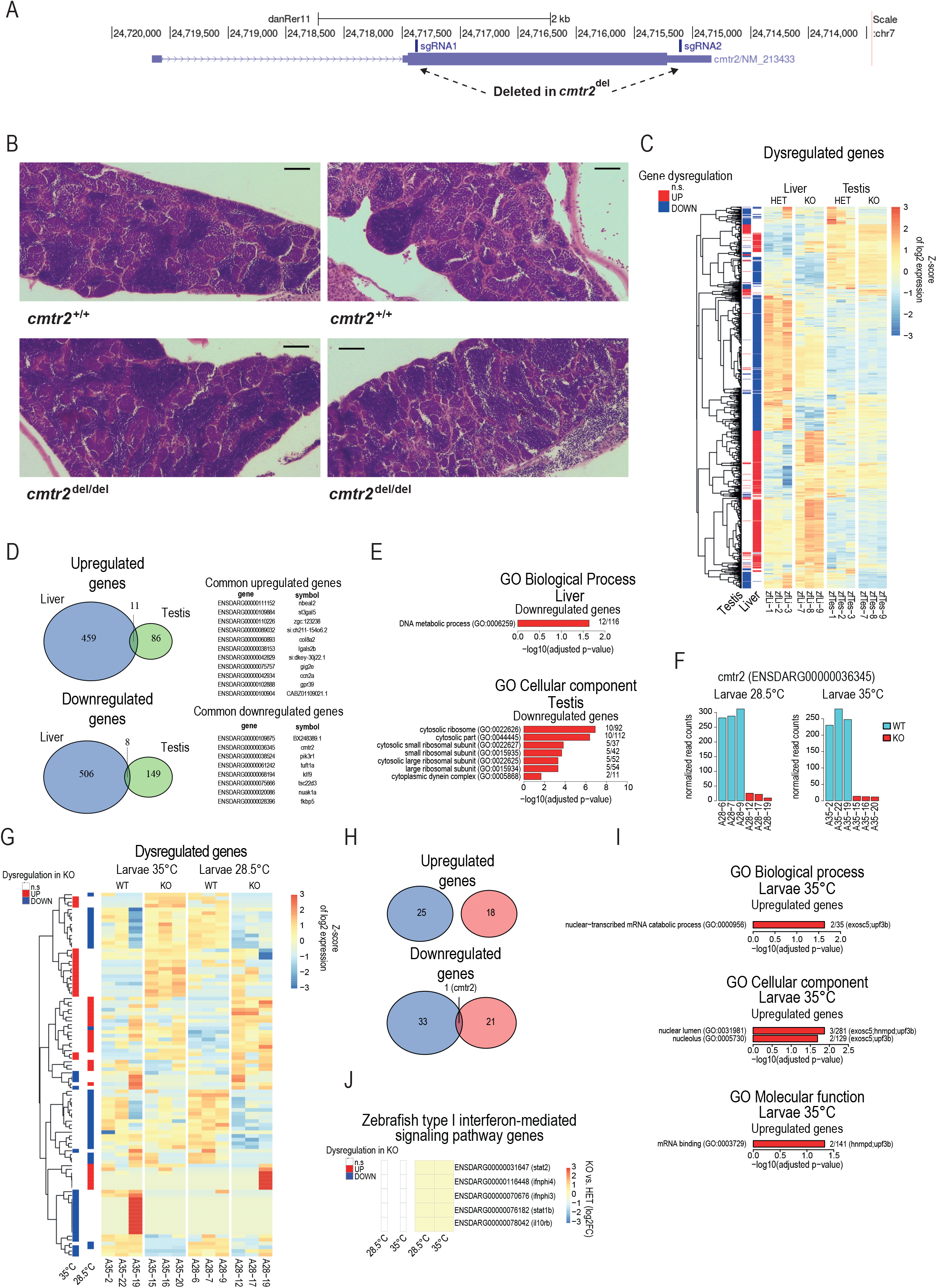
Characterization of gene expression alterations upon *Cmtr2* cKO in zebrafish. (A) Schematic for the generation and initial characterization of *cmtr2* knockout zebrafish. A mutated *cmtr2* allele with a 2.3 kb deletion was generated by targeting *cmtr2* with CRISPR-Cas9 using 2 sgRNAs as shown. The upper part of the panel was adapted from the UCSC Genome Browser. (B) Histological analysis of hematoxylin and eosin-stained sections of Control (*cmtr2^+/+^*) and *cmtr2* KO (*cmtr2^del/del^*) testes. 2 WT and 2 KO fish are shown as examples. Scale bar represents 50 μm. (C) Heatmap showing the expression pattern of all the dysregulated genes in Control (Het) and *cmtr2* cKO liver and testis tissues from adult zebrafish (right). Differentially expressed genes (adjusted p-value < 0.1) are highlighted in red (upregulated) or blue (downregulated) (left). (D) Venn diagrams were generated to determine the common upregulated and downregulated genes in zebrafish tissues upon *cmtr2* knockout. (E) The enriched gene ontology categories for the dysregulated genes in liver and testis are shown. (F) Confirmation of *cmtr2* knockout at RNA level. Expression levels of *cmtr2* RNA in Control and *cmtr2* KO 5 dpf larvae kept at 28.5°C and 35°C. (G) Heatmap showing the expression pattern of all the dysregulated genes in Control and *cmtr2* cKO larvae at the indicated temperatures (right). Differentially expressed genes (adjusted p-value < 0.1) are highlighted in red (upregulated) or blue (downregulated) (left). (H) Venn diagrams were generated to determine the common upregulated and downregulated genes at different temperatures. (I) The gene ontology categories enriched among dysregulated genes upon heat shock. (J) Heatmap showing the expression pattern of Zebrafish type I interferon-mediated signaling pathway genes in zebrafish *cmtr2* cKO tissues as compared to the respective Control tissues at the indicated temperatures (right). Differentially expressed genes (adjusted p-value < 0.1) are highlighted in red (upregulated) or blue (downregulated) (left).

