## Supplementary material for "Cap-specific second nucleotide ribose methylase CMTR2 is required for transcriptome transition during mammalian germline development": Table S1

| Genotyping primers information | | | |
|---|---|---|---|
| **Gene** | **Primer Name** | **Oligo sequence** | |
| ***Cmtr2* KO** | **MD280: KO_fwd** | CCTCCCACACCTCCCCCTGAACCTGAAAC | |
| | **MD282: WT_fwd** | CTACAGAATGTTCATGACCTCTTGAGCAAGC | |
| | **MD283: Common rev** | GTCACAAAACGGCTGCACGTCATCAATGCCG | |
| ***Cmtr2 loxP* 5'end** | **MD 408: Fwd** | GTTTCTAGTTTCCTAGGCAGTTAAGGG | |
| | **MD 409: Rev** | AAGAGGTGGAGCCTAATGGAAGGAA | |
| **Cmtr2 loxP 3'end** | **MD 410: Fwd** | CCTCTGCAAGAACAGCAAATGCTC | |
| | **MD 411: Rev** | CATGTTTATGGCAAGCAAGAAGCAG | |
| **Sftpc CreERT2** | **MD 415: Common fwd** | TGCTTCACAGGGTCGGTAG | |
| | **MD 416:WT rev** | CATTACCTGGGGTAGGACCA | |
| | **MD 414: KI_rev** | ACACCGGCCTTATTCCAAG | |
| **Alb CreERT2** | **51 (MD366), ABV93: Fwd** | GGAACCCAAACTGATGACCA | |
| | **49 (MD365), ABT290: WT Rev** | ATCATTTCTTTGTTTTCAGG | |
| | **50 (MD367), ABT294: KI Rev** | TTAAACAAGCAAAACCAAAT | |
| **DDX4-Cre** | **MM113: Transgene Fwd** | CACGTGCAGCCGTTTAAGCCGCGT | |
| | **MM114: Transgene Rev** | TTCCCATTCTAAACAACACCCTGAA | |
| | **MD 418: Internal ctrl Fwd** | CTAGGCCACAGAATTGAAAGATCT | |
| | **MD 419: Internal ctrl Fwd** | GTAGGTGGAAATTCTAGCATCATCC | |
| ***Cmtr2* N term 3X FLAG-HA** | **HR25_NtermFLAGHA_Fwd** | TGGCTGTCTAGGCCTCTCCCCT | |
| | **HR26_NtermFLAGHA_Rev** | AGAGCCCTGAACTCGAGGTGCC | |
| **PCR amplification primers** | | | |
| **A549-Ace2 KO cell lines validation** | **HR15 Fwd** | AGTAAGTGCAGAAAGACACCAG | |
| | **HR16 Rev** | AACTCCAATGACAAGGAAACCG | |
| **HEK293F KO cell lines validation** | **Fwd** | AGCTTCCTGTGGAATAAGTGGA | |
| | **Rev** | TGCTGCTCATGAGATTTCGC | |
| **Mouse CMTR2 qRT-PCR** | **HR13: Fwd** | CTGTGCAGTTTCCCCCTCAT | |
| | **HR14: Rev** | TCCTGAAGCCCGGTCAGATA | |
| **Mouse β-actin qRT-PCR** | **Fwd** | CCACCATGTACCCAGGCATT | |
| | **Rev** | CGGACTCATCGTACTCCTGC | |
| **Oligos as template for Sensor I (TSS underlined)** | **HR_11_sensor fwd** | TAATACGACTCACTATAGATCTTCCTCTCTCTTCTCCTTCTCCTCTCTTCTCCCTTCTCTCC | |
| | **HR_12_sensor rev** | GGAGAGAAGGGAGAAGAGAGGAGAAGGAGAAGAGAGAGGAAGATCTATAGTGAGTCGTATTA | |
| **Splint DNA used for generating SensorII** | **HR044_Splint** | CTTGGAAGCCATGGTGGCTAGCAAGGGAGAAGAGAGGAGAAG | |
| **Oligos used as templates for synthesis of RNA for *in vitro* methylation (TSS underlined)** | **ELD125: RNA 1 Fwd** | CTCTAATACGACTCACTATTAACCCCCCCCCCCCCCCCCC | RNA 1: m7GpppA-A-CCCCCCCCCCCCCCCCCC |
| | **ELD126: RNA1 Rev** | GGGGGGGGGGGGGGGGGGTTAATAGTGAGTCGTATTAGAG | |
| | **ELD127: RNA2 Fwd** | CTCTAATACGACTCACTATTAGCCCCCCCCCCCCCCCCCC | RNA 2: m7GpppA-G-CCCCCCCCCCCCCCCCCC, |
| | **ELD128: RNA2 Rev** | GGGGGGGGGGGGGGGGGGCTAATAGTGAGTCGTATTAGAG | RNA 9: m7GpppAm-G-CCCCCCCCCCCCCCCCCC |
| | **ELD129: RNA3 Fwd** | CTCTAATACGACTCACTATTACTTTTTTTTTTTTTTTTTT | RNA 3: m7GpppA-C-UUUUUUUUUUUUUUUUUU |
| | **ELD130: RNA3 Rev** | AAAAAAAAAAAAAAAAAAGTAATAGTGAGTCGTATTAGAG | |
| | **ELD131: RNA4 Fwd** | CTCTAATACGACTCACTATTATCCCCCCCCCCCCCCCCCC | RNA 4: m7GpppA-U-CCCCCCCCCCCCCCCCCC |
| | **ELD132: RNA4 Rev** | GGGGGGGGGGGGGGGGGGATAATAGTGAGTCGTATTAGAG | |
| | **ELD133: RNA5 Fwd** | CTCTAATACGACTCACTATAGACCCCCCCCCCCCCCCCCC | RNA 5: m7GpppG-A-CCCCCCCCCCCCCCCCCC |
| | **ELD134: RNA5 Rev** | GGGGGGGGGGGGGGGGGGTCTATAGTGAGTCGTATTAGAG | |
| | **ELD135: RNA6 Fwd** | CTCTAATACGACTCACTATAGGCCCCCCCCCCCCCCCCCC | RNA 6: m7GpppG-G-CCCCCCCCCCCCCCCCCC |
| | **ELD136: RNA6 Rev** | GGGGGGGGGGGGGGGGGGCCTATAGTGAGTCGTATTAGAG | |
| | **ELD137: RNA7 Fwd** | CTCTAATACGACTCACTATAGCTTTTTTTTTTTTTTTTTT | RNA 7: m7GpppG-C-UUUUUUUUUUUUUUUUUU |
| | **ELD138: RNA7 Rev** | AAAAAAAAAAAAAAAAAAGCTATAGTGAGTCGTATTAGAG | |
| | **ELD139: RNA8 Fwd** | CTCTAATACGACTCACTATAGTCCCCCCCCCCCCCCCCCC | RNA 8: m7GpppG-U-CCCCCCCCCCCCCCCCCC |
| | **ELD140: RNA8 Rev** | GGGGGGGGGGGGGGGGGGACTATAGTGAGTCGTATTAGAG | |
